# Cryo-EM structures of insect IRs reveal the structural adaptations of a glutamate receptor ancestor for environmental sensing

**DOI:** 10.64898/2026.09.30.755715

**Authors:** Guido D. Petrovich, Andy Quaen Chen, Marija Simjanoska, Jiawei Zhao, Jaewook Ryu, José Manuel Latorre Estivalis, Marcelo Lorenzo, Josefina Inés del Mármol

## Abstract

Ionotropic receptors (IRs) are an ancient, massively expanded family of sensory ion channels that mediate chemical, thermal, and hygrosensory detection across invertebrates. Derived from synaptic ionotropic glutamate receptors (iGluRs), IRs achieve this diverse tuning by assembling conserved co-receptor subunits with highly divergent tuning subunits into obligate heteromers. To understand how this architecture arose from a glutamate receptor ancestor we report cryo-EM structures of three insect IR complexes spanning the major functional subfamilies: IR8a/IR75b, IR25a/IR76b, and IR25a/IR93a/IR40a. All IR complexes retain the tiered iGluR architecture but adopt a distinctive splayed, non-swapped architecture that our phylogenetic reconstructions relate to a sister branch of glutamate delta (GluD) receptors. Our structures reveal a novel, co-receptor-specific helical ‘beam’ feature that dictates stoichiometry by interlocking the co-receptors and separating the tuning subunits, at the cost of abolishing ligand binding in co-receptors. Comparison of IR8a- and IR25a- containing complexes further illuminates the structural basis of subfamily specificity yielding a predictive code of co-receptor assembly. Lastly, we identify IR-specific coupling mechanisms between the gating machineries of tuning and co-receptor subunits, offering a model for the cooperative gating of IRs. These analyses illuminate how an ancient synaptic receptor was architecturally remodeled into a multimodal family of sensory receptors that drive critical behaviors across invertebrates.

## INTRODUCTION

Organisms perceive the external world using a diverse arsenal of sensory receptors. Chemoreceptors in particular are consistently among the fastest-evolving members in animal genomes (Bargmann 2006, Nei, Niimura & Nozawa 2008): the rigors of a vast and discrete space of ethologically relevant compounds demand a constantly updating repertoire of receptors that detect and distinguish them. A recurrent evolutionary theme to meet this demand is to repurpose internal signaling receptors for environmental sensing. For example, a massive expansion of nicotinic acetylcholine receptors, which typically function in synaptic transmission, underlies the unique ability of cephalopods to “taste by touch” (van Giesen 2020, Kang 2023). A similarly profound evolutionary leap occurred in protostomes, where an ancient family of ionotropic glutamate receptors (iGluRs) was co-opted to form one of the largest and most diverse families of environmental sensors: the invertebrate variant ionotropic receptors, or IRs (Benton 2009, Croset 2010).

IRs boast a long evolutionary history and have undergone massive diversification to detect an impressively large stimulus space that includes chemicals, temperature, and humidity (Benton 2009, Croset 2010, Rytz 2013, Ni 2021). This diversification has produced an unusually elaborate heteromeric logic: they assemble as ligand-gated ion channels that unite broadly expressed co-receptor subunits IR25a and IR8a with stimulus-specific tuning subunits and, in some cases, secondary co-receptor subunits IR76b and IR93a (Abuin 2011, Prieto-Godino 2017, Ni, 2021). In insects, IRs generally fall into three functional groups based on their co-receptor requirements, with each group specializing in distinct sensory modalities.

The IR8a group is primarily expressed in the antenna, the major olfactory organ of insects, where it detects volatile acids, aldehydes, and ketones (Ai 2010, Abuin 2011, Silbering 2011, Prieto-Godino 2017). IR8a-dependent receptors are therefore important for the detection of ecologically relevant odorants, guiding insects towards food sources or away from potentially noxious stimuli (Ai et al. 2010, Abuin et al. 2011, Silbering et al. 2011, Prieto-Godino et al. 2017). In disease-vector mosquitoes, this pathway has been co-opted for host seeking, supporting host localization, blood feeding and, consequently, the transmission of mosquito-borne pathogens (Raji et al. 2019, Cooke 2025). Likewise, this pathway mediates the detection of feeding and oviposition sites in dipteran, lepidopteran, and hemipteran pests that routinely cause agricultural devastation (Zhang 2022, Zhang 2023, Peng 2024, Guo 2023, Shi 2026, Wang 2026). IR8a-containing complexes are therefore high-priority targets for structural determination to enable the development of new chemical strategies for disrupting the life cycles of disease vectors and pests (Ray 2023).

The IR25a group is highly versatile. IR25a/IR76b-containing complexes are found in the antenna and in the mouthparts, mediating detection of amines and diverse taste stimuli from salts and amino acids to carbonation (Silbering 2011, Zhang 2013, Croset 2016, Ganguly 2017, Pitts 2017, Sánchez-Alcañiz 2018, Vulpe 2021). In turn, IR25a/IR93a-containing complexes extend beyond chemosensation transducing thermal and humidity stimuli (Ni 2016, Frank 2017, Knecht 2017, van Giesen 2017, Ni 2021). In disease-vector mosquitoes, IR25a/IR93a/IR21a complexes mediate heat-seeking behavior toward warm hosts (Greppi 2020). IR25a/IR93a-containing complexes in hygrosensory pathways facilitate detection of humidity gradients accompanying the warm, moist boundary layer surrounding a host, and can guide mosquitoes toward suitable oviposition sites (Laursen 2023, Tang 2024, Keane 2026).

IR complexes form the workhorse of environmental sensing across invertebrates, including detection of physical environmental variables that shape host-seeking and reproductive behavior in disease-vectoring insects and agricultural pests. Indeed, ablation of IR-containing complexes leads to the largest behavioral disruption of human host-seeking in the disease-vector *Aedes aegypti*, a behavior that is nearly undisturbed by ablation of the canonical odorant receptor (OR) pathway (DeGennaro 2013, De Obaldía 2022). Furthermore, IRs likely mediate host-seeking sensory processes in arthropods such as ticks and mites (Hoy 2016, Josek 2018, Lei 2019, Zhang 2019, Luciano 2026) that lack insect-specific olfactory pathways. Despite their profound biological importance, the lack of available experimental structures of IRs, coupled to a sparsity of reproducible functional assays, has left a major gap in our understanding of the features of IRs that support host-seeking in disease vectors and, more broadly, stimulus sensing across invertebrates. In this work we initiate the structural characterization of this complex family of receptors by using cryogenic electron microscopy (cryo-EM) to determine the structures of recombinant IR complexes from all three major functional groups: IR8a, IR25a-IR93a and IR25a-IR76b. These structures reveal the structural rules that enforce subfamily specificity, suggest mechanisms by which these channels may be activated, and illuminate the molecular adaptations that allowed an ancient synaptic scaffold to be rebuilt into a versatile platform for environmental sensation.

## RESULTS

We carried out the structural determination of IR complexes representing each of the functional subgroups. We will first focus on the simplest assembly, IR8a/IR75b, to identify defining structural features of the IR family and evaluate its divergence from the parental iGluR branch. We will then examine two IR25a-containing complexes: IR25a/IR93a/IR40a, and IR25a/IR76b, and infer structural features and assembly rules that characterize each IR subfamily.

The IR8a family assembles as heteromers containing the IR8a co-receptor and tuning subunits, without secondary co-receptors. We determined the cryo-EM structure of a heteromeric complex formed by IR8a and IR75b from the kissing bug *Rhodnius prolixus*, a major vector of Chagas disease (Figure 1, Supp. figure 1, 2), in the absence of ligands (apo state).

**Figure 1.**
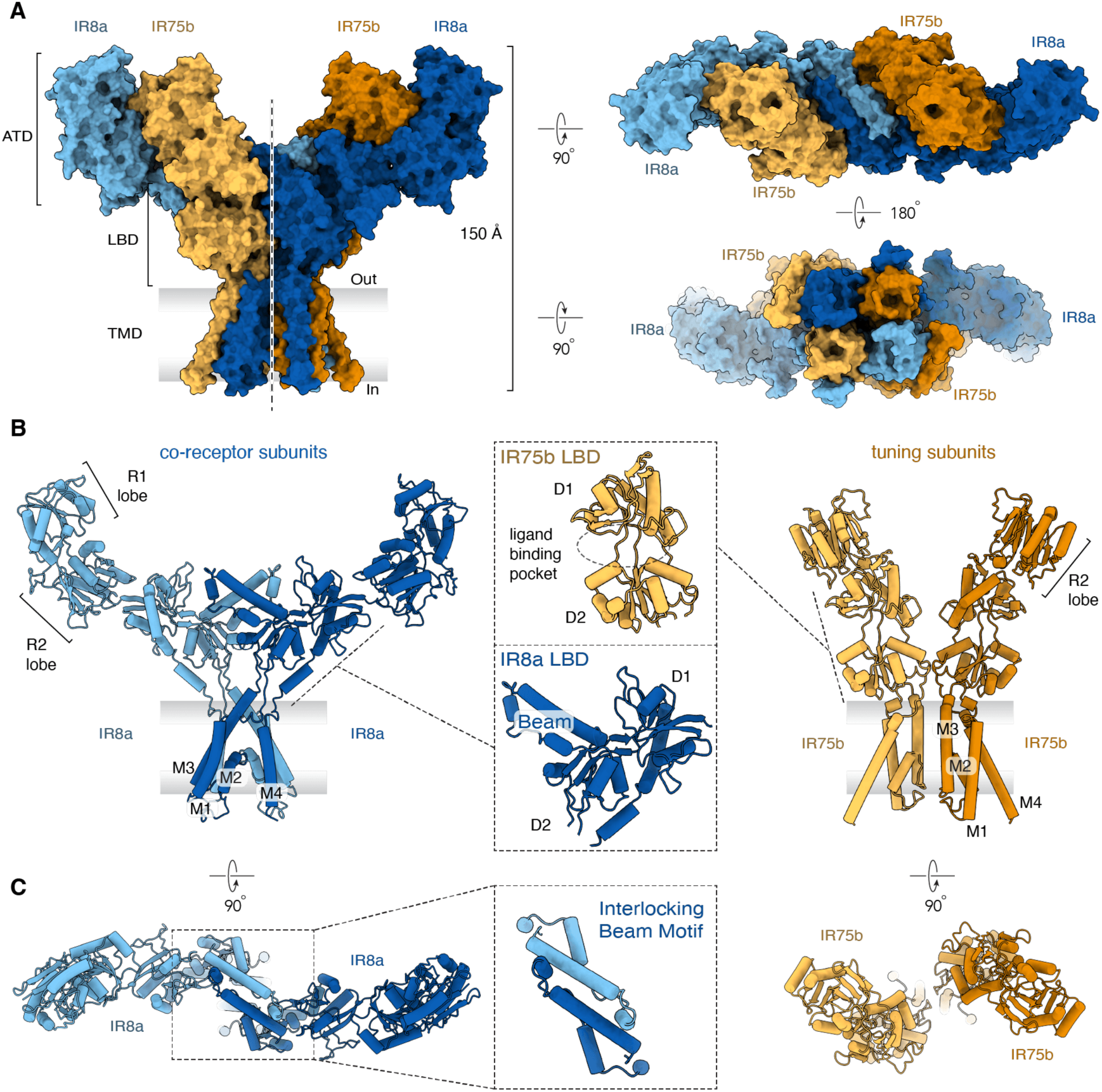
Overall architecture of the RproIR8a/IR75b complex. (A) Surface representation of the RproIR8a/IR75b heterotetramer, colored by subunit (IR8a, blue; IR75b, orange), in side view (left) and in top/bottom views (right), showing the 2:2 stoichiometry, the global two-fold (C2) rotational symmetry, and the wide, T-shaped splay of the amino-terminal domain (ATD) layer relative to the ligand-binding domain (LBD) and transmembrane domain (TMD) layers. (B) Ribbon representation of a co-receptor (IR8a, left) and tuning (IR75b, right) subunit pair. The lobes of the ATD (R1 and R2) and LBD (D1 and D2) domain are marked, noting that IR75b has only the R2 lobe. The TMD M1, M2, M3, and M4 transmembrane helices are labeled. Insets compare the LBD of IR75b, which retains a canonical clamshell fold enclosing a ligand-binding cleft, with that of IR8a, in which the D2 lobe is remodeled and gives rise to the co-receptor beam. (C) Top view of the co-receptor (IR8a) pair, with an inset highlighting the interlocking, three-helix beam motif formed at the interface between the two subunits, and top view of the tuning (IR75b) pair, which makes minimal reciprocal contact.

**Figure 2.**
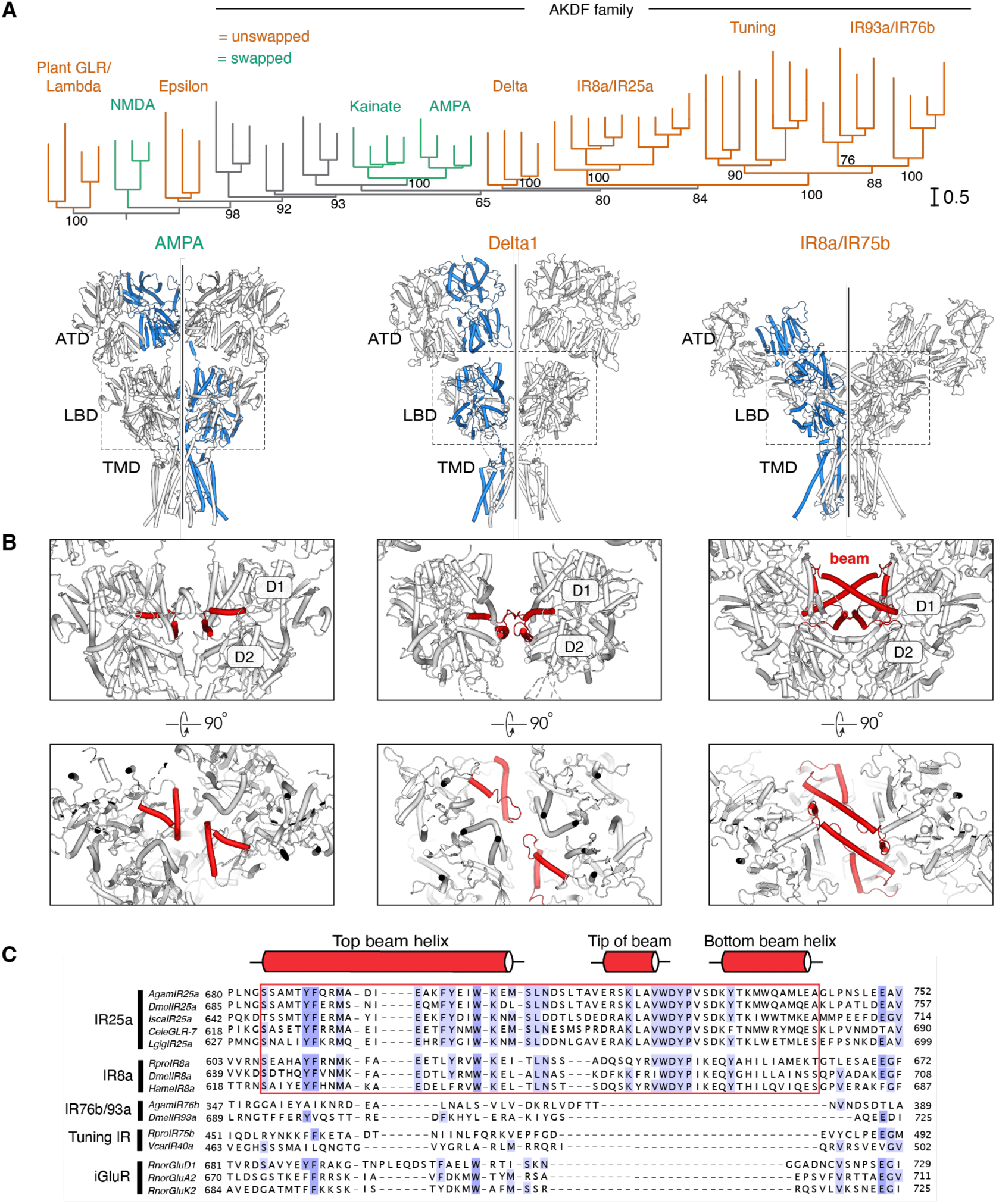
The IR8a/IR75b complex adopts an ancestral, non-domain-swapped iGluR architecture. (A) Maximum-likelihood phylogeny of IRs and other iGluR family members, with domain-swapped (green) and non-domain-swapped (orange) clades indicated. Structures of an AMPA, Delta and this IR8a/IR75b complex are shown with one subunit highlighted in blue, to visualize domain swapping only in AMPA. (B) Top-down views of the LBD layer of the same three receptors, with the D1 and D2 lobes indicated and the region precursor to the beam (red) highlighted, showing its progressive elaboration from a short loop in Delta1 to the extended IR8a/IR75b beam. (C) Sequence alignment of the co-receptor beam helix and adjoining regions across the iGluR and IR family, with conservation shaded.

Overall, RproIR8a/IR75b forms a heterotetramer with a 2:2 stoichiometry, featuring a large extracellular domain, and a global two-fold (C2) rotational symmetry about the vertical axis normal to the membrane. Each half of the complex is formed by a tightly interacting pair of IR8a and IR75b subunits, with identical subunits occupying diagonal positions within the tetrameric assembly (Figure 1a).

The IR8a/IR75b complex displays a distinctive three-tiered architecture that closely resembles that of canonical iGluRs (Sobolevsky 2009, Hansen 2021) (Figure 1a). On the extracellular side are the amino-terminal layer, contributed by the amino-terminal domains (ATD) of each subunit, and the middle layer that joins all four ligand-binding domains (LBD). The transmembrane domain (TMD) that forms the ion channel is contributed by all subunits, each with three fully transmembrane helices (M1, M3 and M4), a re-entrant helix (M2) and pore loop, that in iGluRs, forms the selectivity filter (Figure 1b).

As in iGluRs, the ATDs and LBDs of the IR complex exhibit a bi-lobed clamshell fold, but with marked differences between IR8a and IR75b. While the IR8a ATD is a complete clamshell as in iGluRs, that of IR75b is truncated in half, only including the R2 lobe (Figure 1b). In addition, while the IR75b LBD retains the canonical clamshell shape with a clearly visible cleft, which in iGluRs forms the primary ligand binding site, the IR8a LBD has an additional helical bundle extending from the lower lobe D2 (Figure 1b, c).

Canonical subfamilies of iGluRs are often classified into ‘Y-shaped’ and ‘O-shaped’, determined by the packing and proximity of the ATD domains (Sobolevsky 2009, Herguedas 2016). RproIR8a/IR75b has an exceptionally wide distance between both halves of the ATD layer, resembling a letter ‘T’. Because the ATDs sprawl outwards rather than vertically, the RproIR8a/IR75b complex is much shorter than canonical iGluRs, with a total length of only ∼150Å, compared to ∼170-180Å in iGluRs such as AMPA (Figure 1a, 2a).

### The interlocking beam motif forms a central scaffold at the heart of the complex

Early pioneering studies identified a defining feature in the IR family, found exclusively in the primary co-receptors IR8a and IR25a, and absent in iGluRs and IR tuning subunits (Abuin 2019). This large motif, that extends out of the lower D2 lobe of the co-receptors LBD (Figure 1b, c, 2b, c), is necessary for proper receptor trafficking *in vivo*. It was originally interpreted as an outward-facing loop and dubbed the Co-Receptor Extra Loop (CREL) (Abuin 2019). In the structure of RproIR8a/IR75b, we find this co-receptor motif turned inward, folding into a three-helix bundle that interlocks with the corresponding motif from the diagonal co-receptor to form a prominent structural ‘beam’ at the heart of the complex (Figure 1b, c, 2b). This structural feature is responsible for the striking architecture of this IR complex: it enforces exclusive contacts between the IR8a LBDs, mediates receptor interactions with all other subunits, and causes the ATDs to splay out at an angle (Figure 1, 2).

The beam motif is conserved in all IR co-receptor homologs across protostomes, including in spiralians such as mollusks or annelids that diverged from arthropods >550 million years ago (Croset 2010, Carlisle 2024). Structures of homotetrameric GluD receptors suggest a precursor to the beam: like in IRs, only the two medial-most two GluD LBD subunits contact each other, mediated by a short loop in the corresponding D2 region (Figure 2b, c) (Burada 2019, 2020). However, the exceptional elongation of this motif into the IR beam represents a dramatic evolutionary innovation that likely precluded co-assembly with sister iGluRs, driving the independent diversification of the IR family.

The prominent structural role of the beam is most evident in a top view of the complex (Figure 1a, c, 2b). The two IR8a subunits establish extensive inter-subunit interactions that cross the mid-plane of the complex, totaling 1,414 Å^2^ of buried surface area. In contrast, both IR75b subunits are almost entirely isolated from each other, with only 153 Å^2^ of contact area amongst them, limited to the TMD. Instead, each IR75b subunit is strongly interacting with its adjacent IR8a subunit, with a total of 4,585 Å^2^ buried surface area (Supp. figure 3). Further strengthening the interaction between IR8a and IR75b are two interfacial ion binding sites: a chloride ion pinning the ATDs together and a calcium ion in the LBD apex interface (Supp. figure 10). Both ion-binding sites are present in some iGluRs. The ATD site functions as an anionic pocket across homologous AMPA and Delta receptor interfaces while the latter calcium binding site occurs in a conserved modulatory hotspot for LBD dynamics which recurrently evolved to coordinate different ions, binding calcium in Delta and sodium in Kainate (Elegheert 2016, Naur 2007, Plested 2009, Clayton 2009, Chaudhry 2009a/b, Hansen 2009, Bowie 2010).

**Figure 3.**
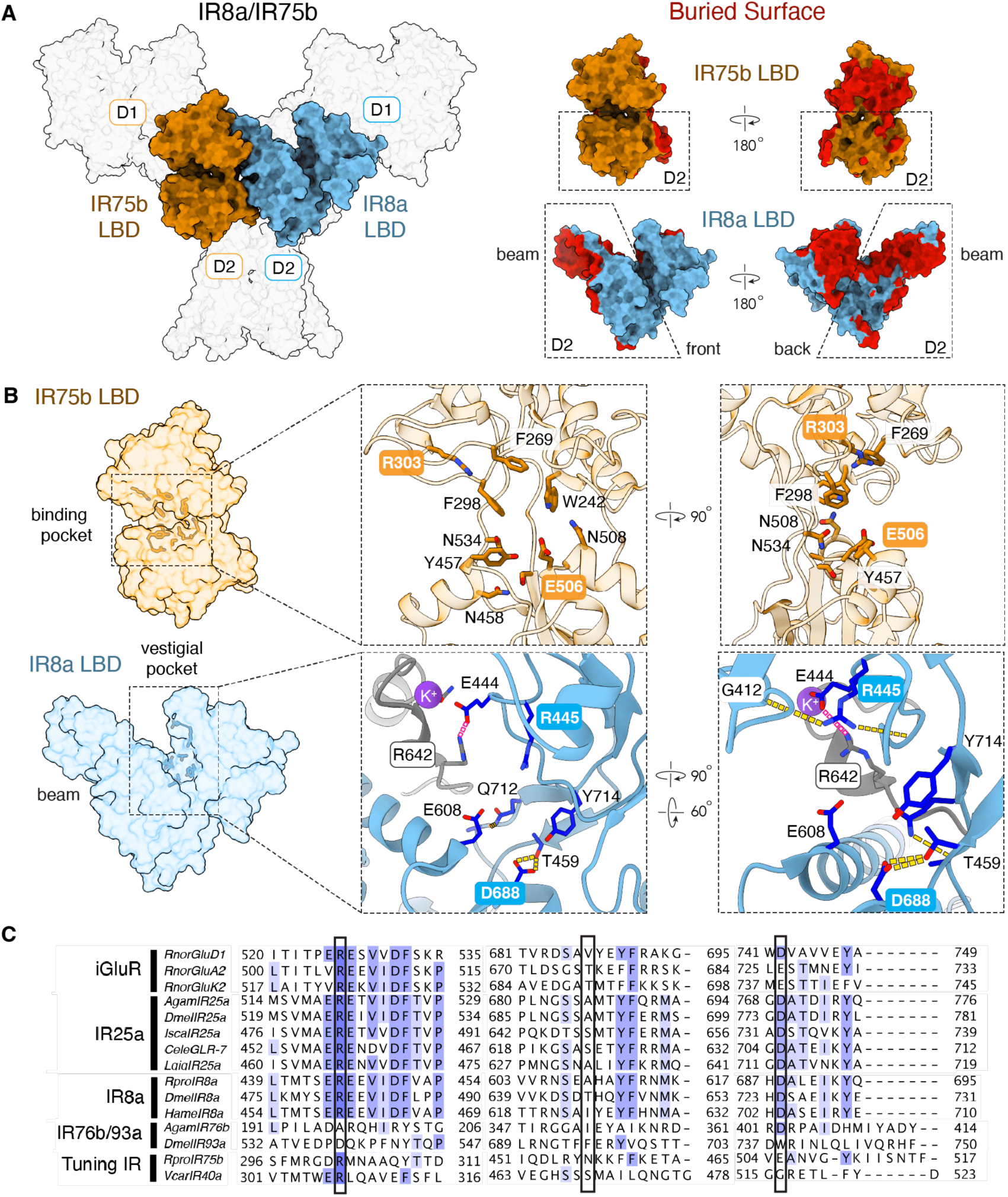
Structural remodeling of the co-receptor LBD locks the IR8a clamshell closed. (A) Surface representation of IR8a/IR75b with the LBD layer highlighted (left), and the isolated LBDs of IR75b and IR8a (right). D1 and D2 lobes are indicated and the inter-subunit buried surface area highlighted in red, showing the extensive contacts made by the IR8a beam relative to the comparatively isolated IR75b LBD. The D2 lobe of IR75b exhibits minimal contacts and is free to move, whereas the extensively remodeled D2 of IR8a is tightly locked in contacts (B) Close-up views of the IR75b (top) and IR8a (bottom) LBD clefts, showing conservation of ancestral glutamate-binding residues (IR75b R303, E506; IR8a R445, D688) and their repurposed structural interactions in IR8a, including the potassium ion (purple) coordinated by beam residue R642 and cleft residue E444. (C) Sequence alignment of the LBD cleft residues across iGluRs, IR25a, IR8a, IR76b/IR93a, and tuning IRs, highlighting conservation of the ancestral glutamate-binding triad.

### An ancestral un-swapped architecture facilitates diversification

Many iGluRs including AMPA, Kainate and NMDA receptors have domain-swapped architectures. In these assemblies, the medial pair of subunits dimerizes with one set of lateral subunits at the level of the LBD, but then crosses over to interact with the other lateral subunits at the ATD tier (Figure 2a). This domain organization results in a tightly packed and highly interlocked subunit architecture, extensively connecting all subunits with one another (Sobolevsky 2009, Das 2010, Karakas 2014, Lee 2014). In contrast, RproIR8a/IR75b has a non-domain-swapped architecture where all domains of each subunit consistently occupy the same relative position to the domains of other subunits. At both the ATD and LBD tiers, each subunit retains the same neighbor, in contrast to the domain-swapped AMPA receptors (for example, PDB:3KG2), where a subunit’s ATD and LBD partners differ (Figure 2a).

This unswapped architecture likely reflects the ancestral form of the iGluR family. We generated Maximum-likelihood (ML) phylogenetic trees containing IRs and other iGluRs (Figure 2a, Supp. files), with dense cross-clade sampling of AKDF (AMPA, Kainate, Delta and Phi) subgroup members which IRs have known affinities with (Croset). We find that IRs form a monophyletic clade (UFBoot2 Boostrap support 84) sister to Delta receptors (BS 80) and squarely within the broader AKDF radiation (BS 93). Recent structures of the sister branch of Delta receptors, as well as outgroup plant GLRs, some NMDAR conformations, and predicted structures of placozoan Epsilon receptors revealed unswapped arrangements (Figure 2a) (Green 2021, Jalali-Yazdi 2018, Burada 2019, 2020, Tian 2021, Singh 2025, Wang 2025).

Structurally, the ancestral unswapped architecture inherently accommodates the addition of the beam motif, establishing the foundation for the complex heteromeric logic that defines the IR family. Insertion of the beam into the medial co-receptor subunits strengthens their interactions, shifting the burden of tetrameric assembly onto the highly conserved co-receptor subunits.

### Structural remodeling of the co-receptor LBDs

As in iGluRs, the IR LBD is a large extracellular domain, physically separated from the transmembrane ion channel domain and connected to it via linkers (Figure 3a). Apo structures of iGluRs generally present open, hydrated LBD clefts containing solvent-exposed, charged residues on both upper (D1) and lower (D2) lobes. The charged iGluR ligands, generally zwitterionic amino acids, satisfy these charges and bridge the cleft, closing the clamshell. This exerts tension on a linker that connects to the ion channel and transmits conformational changes from the LBD to the channel pore (Armstrong 2000; Sobolevsky 2009; Dürr 2014; Twomey 2017, Hansen 2021). Structurally, the IR75b LBD appears to be capable of executing a similar mechanism. Its D2 lobe is minimally entangled in intersubunit interactions and its clamshell therefore free to close when a cognate ligand stabilizes a closed-cleft conformation (Figure 3a). In contrast, the IR8a LBD is extensively remodeled, forced into a sharp vertical angle by the beam domain that ‘lifts’ the lower lobe (D2) upwards and establishes extensive intersubunit interactions with the beam of the diagonal IR8a subunit (Figure 3a). As these inter-beam interactions are central to the tetrameric integrity of the complex, the IR8a LBDs appear to be structurally locked and incapable of executing the clamshell closure mechanism that gates canonical iGluRs.

Examination of LBD clefts in our structure reveals partial conservation of the amino acids that in iGluRs form the canonical glutamate-binding triad (Figure 3 b, c). The most critical residue, an arginine that coordinates the glutamate alpha carboxylate, is conserved in both IR8a (R445) and IR75b (R303). In addition, an acidic residue that coordinates the glutamate alpha amine is also conserved in both the IR8a (D688) and IR75b (E506) (Figure 3b, c; Naur 2007, Mayer 2011, Abuin 2011, Hansen 2021). While the cognate ligand of many tuning subunits (including RproIR75b) is unknown, the IR8a subfamily is generally tuned towards carboxylic acids (Prieto-Godino 2017, 2021). Therefore, the conservation of R303 in IR75b can be well-explained as it could easily coordinate carboxylic acid ligands in a pose similar to that of iGluR ligands to engage the same LBD activation mechanism.

In IR8a, the conservation of ancestral ligand-binding residues presents an interesting puzzle. As a stimulus-inert co-receptor with a fixed LBD, IR8a is not expected to bind activating ligands. Why, then, are these residues conserved? Our structure shows that they have been repurposed to serve a structural role and are permanently engaged in stabilizing interactions within the LBD. R445 establishes critical interactions with T440 and G412 that serve as a two-point anchor of a D1 loop, and D688 forms hydrogen bonds with T459 in the pocket (Figure 3b). Additional stabilization is provided by the beam of the opposing IR8a subunit that wedges its tip directly into the cleft. This insertion is stabilized by a salt bridge between R642 in the beam with E444 in the cleft, jointly coordinating a potassium ion (Figure 3b). Collectively, this dense network of intra- and inter-subunit interactions permanently tethers the cleft of IR8a, supporting that it represents a vestigialized ligand-binding pocket. The structural role of these conserved residues in the IR8a cleft might explain why mutating equivalent *Drosophila* IR8a residues resulted in loss-of-function phenotypes to various degrees *in vivo* (Abuin 2011), a finding that was originally interpreted as suggesting potential ligand-binding activity by IR8a.

### Mechanical coupling of tuning and co-receptor subunits at the pore gate

The TMD is the final effector of receptor-wide conformational changes, directly controlling ion flux and downstream neuronal activity in response to stimulation. Because the central beam occludes the vertical axis of the ectodomain, extracellular ions enter the RproIR8a/IR75b TMD ion conduction pathway through lateral entry tunnels between the D2 domains of each LBD dimer (Figure 1a).

In iGluRs, the gate is formed where the transmembrane M3 helices converge at the extracellular entrance to the pore. Ligand binding stabilizes a closed LBD clamshell conformation by upward swinging of the D2 lobe. This movement exerts tension on a short M3-D2 linker, causing a conserved ‘SYTANLAAF’ sequence in the stalk of the M3 helices to hinge outward, yielding an open pore.

In RproIR8a/IR75b, which is an apo structure, the gate is closed. The narrowest constriction (3.4Å wide) is formed by Q431 at the tips of the IR75b M3 helices (Figure 4a). The IR8a M3 helices rest in a slightly more permissive conformation, separated by a 7Å distance. Since our LBD analyses suggest that only the two tuning receptor subunits are able to undergo LBD clamshell closure in response to ligand binding, it is possible that opening of the two IR75b M3 helices suffices to yield an open state, without major structural rearrangements of the IR8a M3 helices.

**Figure 4.**
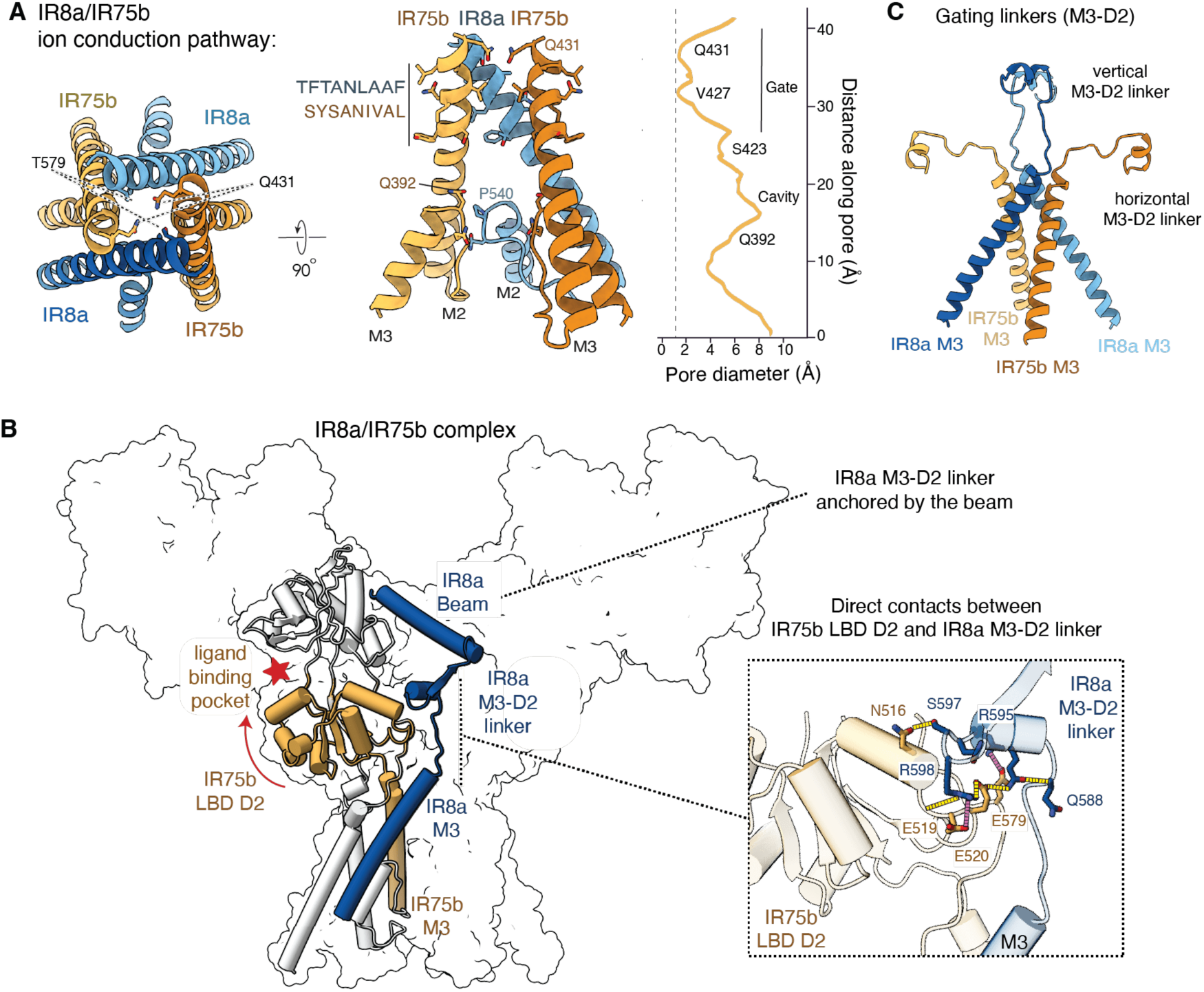
Mechanical coupling between tuning and co-receptor subunits at the pore gate. (A) Top and side views of the pore-lining M3 helices of RproIR8a/IR75b, with the SYTANLAAF-equivalent motifs of IR8a and IR75b indicated and the narrowest constrictions (IR75b Q431) labeled; pore radius plotted as a function of distance along the pore. (B) The unique network of contacts between the IR75b LBD D2 lobe and the IR8a gating M3-D2 linker, with the IR75b ligand-binding pocket indicated and the interacting residues shown in the inset. Arrow shows direction of movement of the D2 lobe upon ligand binding in canonical iGluRs. (C) Comparison of the distal (IR75b) and proximal (IR8a) linker geometries.

However, two structural features suggest that the two IR8a M3 helices might also experience structural rearrangements to yield an open state. First, the SYTANLAAF motif is retained in the co-receptor IR8a subunits (TFTANLAAF), and yet largely degenerated in the tuning IR75b subunits (Figure 4a). Second, the D2 lobe of IR75b makes a unique and extensive network of direct contacts with the M3-D2 linkers of IR8a (Figure 4b), interactions that are entirely absent in iGluRs. This interaction may enable cooperative pore opening of all M3 helices by mechanically coupling the upward swing of the tuning LBD to the M3-D2 linker of coreceptors. Future studies of active states will illuminate the extent of rearrangements across the tetramer that result in an open pore.

Deeper in the transmembrane ion conduction pathway, a second notable constriction is formed by the apex of the M2 re-entrant pore loop, marked by Q392 in IR75b (Figure 4a). In canonical iGluRs, this region houses the Q/R/N selectivity filter, which varies in the iGluR family to tightly control calcium permeability (Hume 1991, Burnashev 1992, Hansen 2021). Sequence alignments of IRs indicate that they lack these canonical residues (Supp. figure 9, Supp. files; Abuin 2011). Instead, this position is generally occupied by a proline in the co-receptors IR8a and IR25a, and by wildly divergent, often hydrophobic residues in secondary co-receptors and tuning subunits. Unlike iGluRs, where the Q/R/N site creates a formidable secondary constriction, our apo RproIR8a/IR75b structure reveals a much wider passage at the M2 apex, nearly 10Å in diameter (Figure 4a). This suggests that the M2 loop may not serve as a stringent selectivity filter in the IRs investigated here.

### Theme and variations: structures of 25a-containing complexes showcase structural diversity within a common general motif

Over its long evolutionary history, the IR family acquired an unusually complex heteromeric logic involving primary co-receptors, secondary co-receptors and tuning receptors. While the primary co-receptor IR8a forms strictly di-heteromeric complexes with tuning subunits, its paralog IR25a often co-assembles with both a tuning subunit and a secondary co-receptor (IR93a or IR76b). These secondary co-receptors phylogenetically cluster among and are likely derived from tuning receptors (Supp. Files, Figure 2a; Croset 2010), and similarly lack the beam domain. Secondary co-receptors also differ extracellularly: IR93a retains a full ATD whereas IR76b lacks one entirely. Functionally, IR25a and IR93a co-assemble with conserved thermo- or hygrosensory tuning subunits (IR21a, IR40a and IR68a). In contrast, IR25a and IR76b partner with a larger array of divergent tuning to detect chemosensory cues like salts and amines.

To characterize the structural basis of subfamily specificity, we elucidated the cryo-EM structure of a complex containing IR25a/IR93a/IR40a, a “dry receptor” necessary for hygrosensation (Supp. figure 4, 5, 6, 7) (Frank 2017, Knecht 2017). To maximize biochemical stability, we used a chimeric strategy that has been successful in other fields (Silbering 2016, Zhao 2024), combining orthologs from diverse insect species: *Anopheles gambiae* IR25a, *Drosophila melanogaster* IR93a and *Vanessa cardui* IR40a (Lohse 2021).

**Figure 5.**
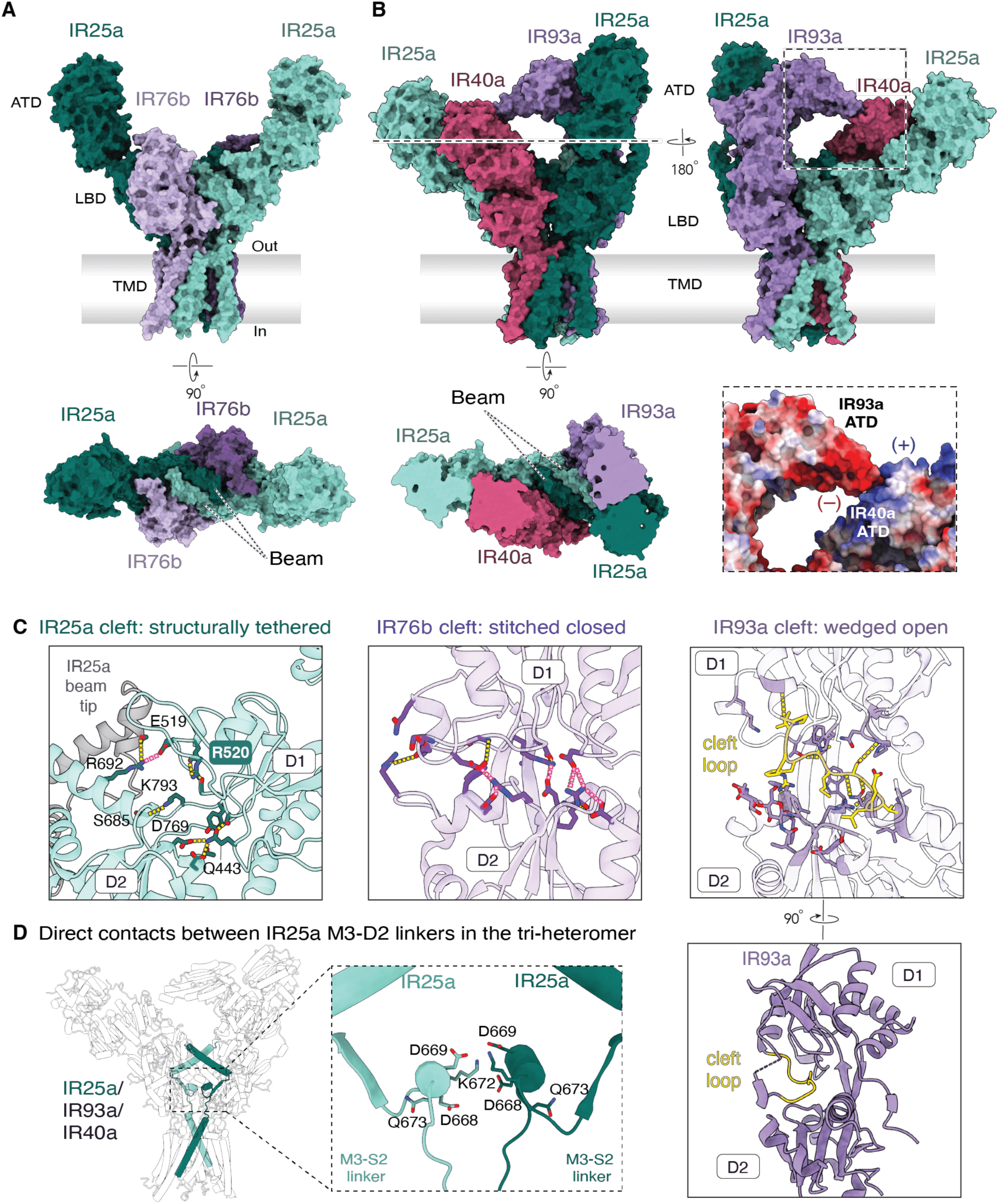
Overall architecture and ATD diversity of IR25a-containing complexes. (A) Surface representation of the AgamIR25a/AgamIR76b complex in side view (top) and top view (bottom), colored by subunit. (B) Surface representation of the asymmetric AgamIR25a/DmelIR93a/VcarIR40a complex in side views (top) and a top view taken at the position indicated by the dashed line (bottom). Inset shows a close-up of the electrostatic interaction that docks the snout of the IR93a ATD in the IR40a ATD. (C) Close-up views of the IR25a (left), IR76b (middle), and IR93a (right) LBD clefts, showing the structurally tethered, closed, and wedged-open conformations, respectively. (D) Unique contacts between the M3-D2 linkers of the two diagonal IR25a subunits in the IR25a/IR93a/IR40a complex (left, overall view; right, close-up), showing the extended coupling that is absent in IR8a and iGluRs.

We also targeted determination of *A. gambiae* IR25a/IR76b/IR41a, which responds to nitrogen heterocyclics and is implicated in oviposition site selection (Pitts 2017, Raji 2023). However, although we confirmed the presence of all three subunits throughout the purification, cryo-EM analysis yielded a high-resolution map of only the IR25a/IR76b di-heterotetramer, with no resolvable density attributable to IR41a (Supp. figure 4, 5, 6). Because IR25a/IR76b are thought to require a tuning subunit for physiological function, it is unclear if this structure represents a functional state. We therefore restricted our analysis of this complex to the interfaces and general architecture of IR76b.

### Overall architecture and subunit composition of IR25a assemblies

Like the IR8a complex, IR25a assemblies retain the tiered ATD, LBD and TMD architecture of ancestral iGluRs (Figure 5a). The IR25a/76b complex forms a 2:2 di-heterotetramer, and the IR25a/93a/40a complex forms a 2:1:1 tri-heterotetramer where IR93a occupies the structural position of a second tuning receptor. Analogously to IR8a, in both IR25a complexes the two IR25a subunits sit diagonally across the central axis, tightly interlocked via their beam domains. As in IR8a, this central beam mediates contacts with all subunits prohibits direct LBD-LBD interactions between the tuning or secondary coreceptor subunits (Figure 5a). The structures of these complexes differ most distinctively at the ATD layer due to the varying lengths of the secondary co-receptor ATDs. In the IR25a/IR76b complex, the ATD layer is composed entirely of IR25a, whose ATDs are loosely packed against the LBDs and extend vertically (Figure 5a). In contrast, the full-sized IR93a ATD forms a ‘bridge’ connecting the ATDs of IR25a and IR40a. This interaction is enabled by the unique ATD of IR93a, containing an elongated helical ‘snout’ capped by an electronegative surface that directly engages two arginine residues in the IR40a ATD (Figure 5a, b). Further stabilizing this unique ATD orientation is an elaborate glycosylation that originates from N475 in the IR93a LBD (D1 lobe) and projects vertically, reaching its own ATD. As a result, the tri-heteromeric ectodomain is highly asymmetric, reminiscent of recent structures of GluD2 receptors (Burada 2020, Wang 2025).

Conserved interfacial ion binding sites strengthen these assemblies. As in the IR8a complex, a chloride ion mediates ATD-ATD interactions between IR25a/IR40 (Supp. figure 10). Similarly, the ancestral LBD cation binding site is preserved in these assemblies, formed by equivalent residues as in iGluRs and the IR8a complex, and coordinating a sodium ion. Notably, the IR25a/IR93 interfaces lack both ion sites. In the LBD layer, the cleft of the IR25a clamshell remains open and lacks ligand density. As in IR8a, the ancestral glutamate-binding residues R520 and D769 establish stabilizing interactions with neighboring D1 loops (Figure 5c) tethered the IR25a cleft in place. In contrast, the LBDs of the secondary co-receptors are more drastically remodeled to preclude ligand binding. In IR76b, the clamshell is closed (Figure 5a, c) despite the absence of any ligand-like densities. Instead, the two lobes have been stitched together by a dense interaction network in which many formerly unsatisfied ligand-binding ionic contacts are conserved in position but supplied with novel partners (Figure 5c). The binding pocket of IR93a is effaced more drastically: an elongated D1 segment is turned into an auto-inhibitory loop, permanently wedged into the cleft through electrostatic and hydrophobic interactions (Figure 5a, c). This, like in IR76b, probably forecloses any opportunity for ligand binding and clamshell movement. However, unlike IR76b, the IR93a clamshell is immobilized in an open rather than closed conformation.

The central pore in the TMD of the IR25a/IR93a/IR40a structure is in a non-conductive state. However, unlike in IR8a/IR75b, where the tuning subunits form the narrowest point, the constriction of the IR25 complex is formed by the two IR25a T656 in the stalk of the M3 helices (Supp. figure 7). IR93a and IR40a rest in a more permissive distance. While this complex is known to mediate humidity sensing, the biophysical stimulus that effectively opens the channel is yet to be determined. Therefore, what conformational changes of the tuning subunit might underlie channel activation is unknown. However, we observe in the IR25a complex an additional set of interactions that might be involved in gating: the M3-D2 linkers of the IR25a are not only contacting the LBDs of neighboring subunits, but also the corresponding M3-D2 linker of the diagonal IR25a subunit across the central axis (Figure 5d). This extended coupling, absent in IR8a and iGluRs, may enable the sole tuning subunit to gate the pore constricted by the co-receptor subunits.

### The beam dictates heteromeric stoichiometry

The presence of the beam only in primary co-receptors offers a structural rationale for the observed stoichiometries of IR complexes. The central, interlocking nature of this feature suggests that exactly two beam-containing subunits need to occupy diagonal positions within the tetramer and likely precludes the incorporation of a third primary co-receptor. Because all tuning IRs and secondary co-receptors lack this domain, our structural data suggests that all IR complexes will likely contain two (and only two) primary co-receptor subunits.

Previous *Drosophila* work identified a conserved N-glycosylation site within the beam of IR8a and IR25a that promoted ciliary localization in a subset of IR8a-dependent receptor complexes (Abuin 2019). We observe clear densities indicating glycans on corresponding residues (IR8a N633, IR25a N710) in all three structures (Supp. figure 2, 5). Curiously, these glycans are clearly oriented differently in IR8a and IR25a-containing complexes. In our IR8a/IR75b structure, the glycan on IR8a N633 directly contacts IR75b E255 in the tuning subunit. The equivalent glycosylated residue in IR25a N710 projects in the opposite direction, and instead interacts with E717 on the beam tip in the same IR25a subunit. Although the exact size and composition of this glycan in our heterologous structures may not represent those in native complexes, it appears to be a versatile feature that can engage in variable stabilizing interactions depending on geometry and availability of local partners, explaining its uneven requirement *in vivo*.

Further examination of the beam architecture suggests that IR25a and IR8a self-associate via distinct mechanisms. In IR8a, a conserved leucine on the top helix (L623) and an isoleucine on the bottom helix (I648) mediate tight hydrophobic packing between the isologous faces of the two beam domains. In IR25a, the equivalent positions are occupied by F700 on the top helix, which engages in isologous π–π stacking, and by V728 on the bottom helix (Figure 6a). These alternative interfaces likely favor homotypic self-association within each co-receptor and disfavor mixed IR8a–IR25a beam pairing. This observation might explain in vivo evidence indicating that IR25a and IR8a do not co-assemble in the same complex, despite sharing overall architectural similarities and being often co-expressed in the same olfactory sensory neurons (Abuin 2011, Ai 2013, Task 2022).

**Figure 6.**
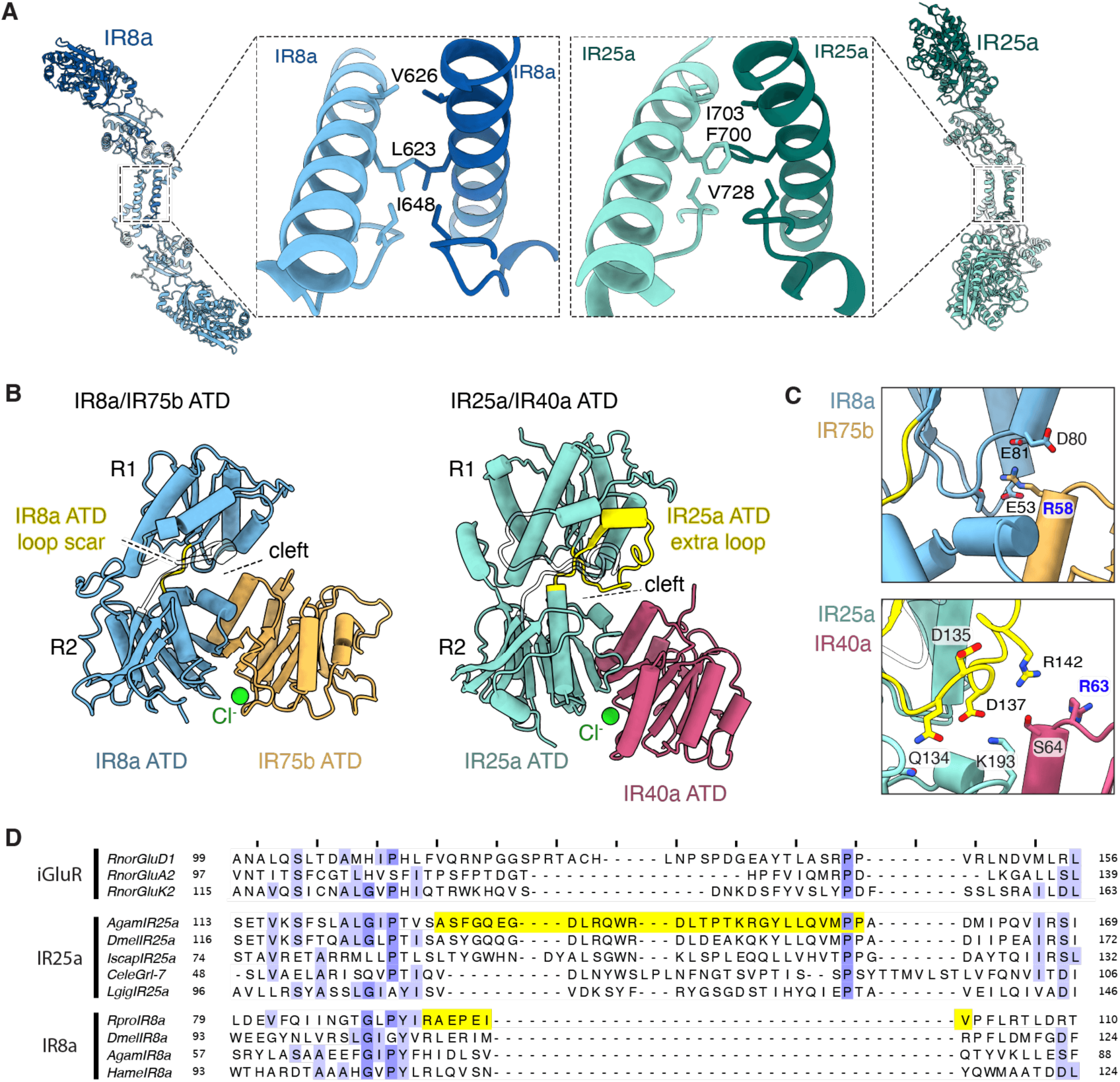
Distinct self-association and ATD-interface mechanisms distinguish IR8a and IR25a. (A) Close-up of the beam-beam interface in IR8a (left) and IR25a (right), showing the residues that mediate homotypic self-association (IR8a L623, I648; IR25a F700, V728). (B) Comparison of the IR8a and IR25a ATD clefts, showing loss of the ancestral ATD loop (yellow) in IR8a relative to IR25a, and the chloride ion (green) at each ATD-ATD interface. (C) Cleft details of the IR8a/IR75b (top) and IR25a/IR93a/IR40a (bottom) complex, highlighting the interactions between the IR40a ATD and the open cleft of IR8a, which are occluded by the conserved loop of the IR25a subunit. (D) Sequence alignment of the ATD interface loop region across iGluRs, IR25a, and IR8a lineages, showing conservation of the ancestral loop in IR25a and its loss in IR8a.

### The structural basis of subfamily specificity

We next sought to identify the structural principles that characterize how IR8a and IR25a recruit specific tuning and secondary co-receptors. We first generated subfamily-specific alignments of tuning subunits with documented in vivo evidence of selective functional association with either IR8a or IR25a (Supp. table 3, files). We then mapped the resulting sequence or structural conservation of each subfamily onto our experimental structures to identify the structural code of subfamily specificity (Katoh 2013, Celniker 2013, Kim 2025).

#### Differential interactions at the ATD interface

Although the ATDs of IR8a and IR25a share an overall similar architecture, they engage their tuning ATDs through different interfacial interactions. This divergence stems from evolutionary variation in the co-receptor clefts (Figure 6b). A a highly hydrophilic and structured loop in the IR25a ATD is wedged into its own cleft and provides internal countercharges to satisfy the cleft’s electrostatic surface. This loop sterically occludes the cleft, pushing the IR40a ATD downward and forcing a shallower interaction footprint (Figure 6c). This loop is of ancestral origin: it is present in delta and kainate receptors and conserved across IR25a (Figure 6d). IR8a has lost this ancestral ATD loop and exposes its strongly electronegative cleft towards the interface with tuning ATDs (Figure 6b, c, d).

The ATDs of tuning subunits mirror these differences: IR75b exhibits a protruding positively charged arginine (R58) reaching directly into the electronegative IR8a ATD cleft (Figure 6c). In contrast, IR25a-dependent tuning subunits (including IR93a) present acidic or hydrophobic residues in the homologous region (Figure 6c).

#### Differential interactions at the LBD interface

In tuning subunits, an L-shaped helix runs along the side of the subunit’s D1 lobe (Figure 7a). In IR25a-dependent tuning subunits (like IR40a), the lower ‘forearm’ helix is longer (I569 to R586), running parallel to the bottom helix of the IR25a beam, and guiding its orientation (Figure 7b). In contrast, in IR8a-dependent tuning subunits (like IR75b), this forearm helix is substantially shortened (L563 to W571) and does not reach the IR8a beam.

**Figure 7.**
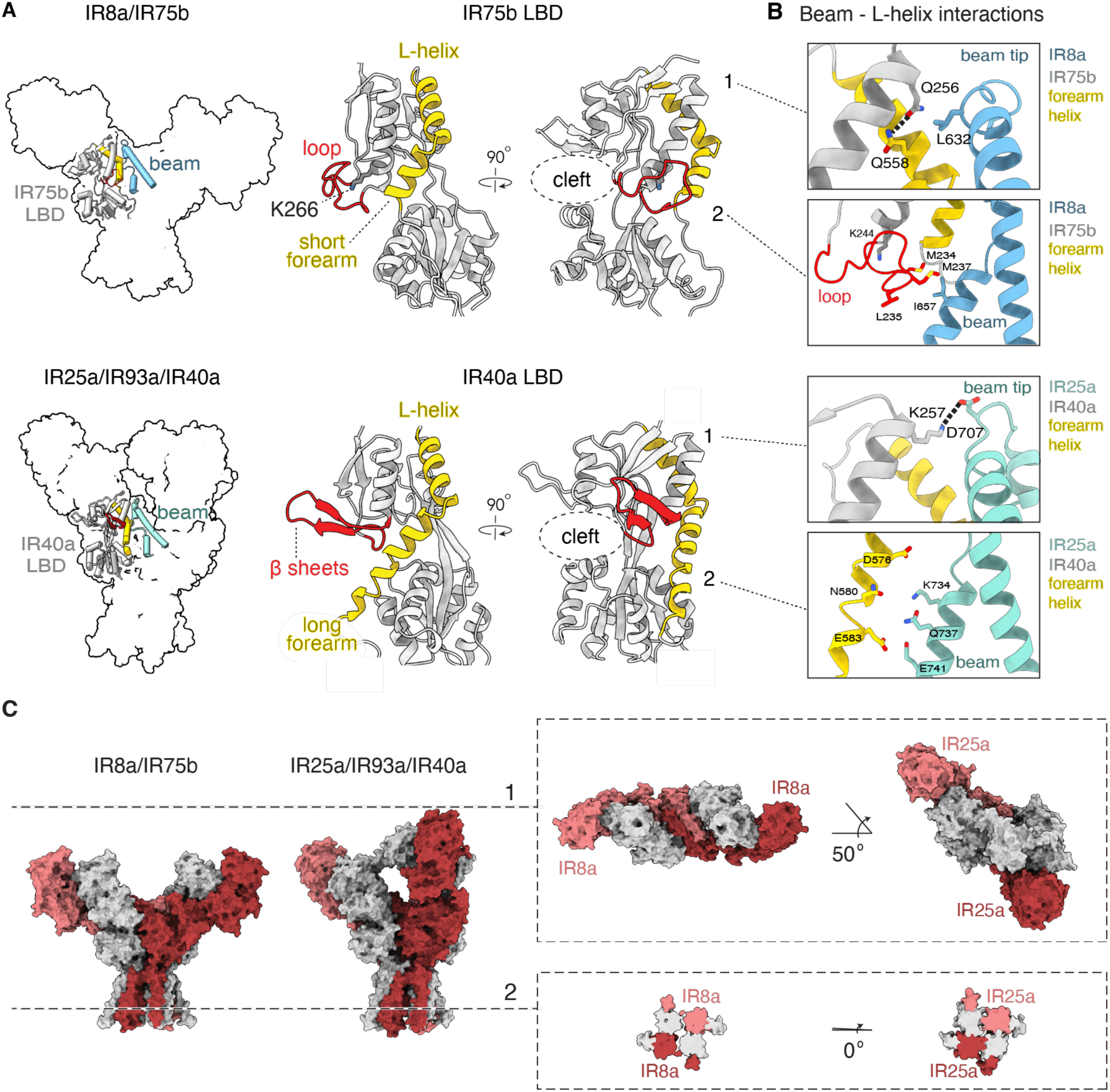
Structural determinants of tuning-subunit subfamily specificity. (A) Left, surface representation of the complexes, highlighting the position of the LBD regions detailed in this figure. Middle, the LBDs of the tuning subunits showing the L-helix, with the forearm region of variable length and presence of a diagnostic loop or beta sheet element. (B) Close-up views of the beam - L-helix interface in IR8a/IR75b (top) and IR25a/IR40a (bottom), showing the distinct anchoring interactions at the top and bottom of the L-helix that stabilize each pairing. (C) Structural alignment of the IR8a/IR75b and IR25a/IR93a/IR40a by the TMD layer shows the global rotation of the ectodomain of the IR25a complex relative to the ectodomain of the IR8a complex.

Adjacent to this extended forearm helix, IR40a features a conserved, prominently structured, outward-protruding two-stranded antiparallel beta sheet bundle (Figure 7a). This ancestral beta sheet bundle is absent in IR75b, and instead, the analogous region forms a conserved structured hydrophobic loop that reaches up to interface with the characteristically hydrophobic bottom helix of the IR8a beam (I657). This loop is rigidly anchored in place by a basic residue (IR75b K244) that is the single most conserved residue across all IR8a-dependent tuning receptors (Figure 7b).

These localized geometric differences in the lower LBD might contribute to a substantial divergence in the global architecture of the two complexes. Structural alignment of the two complexes reveals that the subunit position and topology remain highly conserved in the TMD layer, but diverge markedly in the ectodomain. In IR25a/IR93a/IR40a, the entire extracellular assembly is rotated by ∼50 degree relative to that of the IR8a/IR75b complex (Figure 7c). This twist is likely dictated by the differential packing of tuning subunits against their co-receptor beams.

Further up, conserved contacts at the beam tip reinforce differential interactions. At the IR25a beam tip, a highly conserved acidic residue (D711) forms a salt bridge with a conserved basic residue on the tuning subunit (K257 in IR40a) (Figure 7b). IR8a, with a shorter beam tip, has lost the equivalent interaction. Instead, IR75b forms an intrasubunit hydrogen bond between two highly conserved residues, Q256 and Q558.

### A predictive structural code for IR assembly

Analysis of the conservation of these architectural hallmarks allow us to propose a set of predictive criteria for determining whether an orphan tuning receptor will assemble with IR8a or IR25a.

Assembly with IR8a is characterized by: (1) a short forearm helix (<12 amino acids, found in 100% of IR8a-dependent subunits); (2) a signature basic residue (equivalent to IR75b K244) stabilizing a hydrophobic loop that packs against the beam base (100% conservation); and (3) a positively charged ATD surface at the interface (68% conservation).

Assembly with IR25a is characterized by: (1) an extended forearm helix (>12 amino acids, found in 97% of IR25a-dependent receptors); (2) an outward-protruding, two-stranded beta sheet bundle on the D1 lobe (97% conservation) and (3) a neutral or negatively charged ATD (64% conservation).

## DISCUSSION

In this study, we determined structures of insect IR complexes that sense widely diverse stimuli and incorporate varied heteromeric architectures. A defining feature of this family is the emergence of inert primary co-receptors with extensively remodeled LBDs featuring an interlocking central beam. This structural innovation distinguishes IRs from their ancestral iGluR: rather than forming extensive interactions amongst all LBD subunits as seen in canonical iGluRs, the beam tightly locks the co-receptor LBDs together diagonally. This isolates the tuning subunits and constrains them to only interact with neighboring co-receptors.

The profusion of greatly diversified and mutually incompatible complex types incorporating multiple co-receptor classes in IRs over 550 million years of divergence is effectively a natural evolutionary experiment in molecular assembly and diversification. In this discussion, we will hazard some evolutionary speculations with broader implications based on our structures, with an eye to sparking other inquisitions into how chance and necessity shapes evolution of multimeric assemblies and sensory perception.

### Co-receptors support evolvability of expanded multimeric receptor families

It is curious and noteworthy that obligately multimeric chemoreceptors—generally ion channels, which require multiple pore-forming subunits—often settle in systems in which conserved, broadly expressed co-receptors heteromerize with divergent tuning receptors. The insect OR family, another class of invertebrate chemoreceptors unrelated to IRs, also has independently evolved an obligate heteromeric system where diversified tuning receptors assemble with the co-receptor Orco. Like IRs in protostomes, this heteromeric system emerged before larger expansions of the protein family occurred in insects; no comparable homomeric systems of such magnitude are known so far. Heteromerization partitions ancestral receptor functions between the two subunit classes: coreceptors primarily mediate assembly and complex stability, while tuning receptors provide ligand specificity. Consistent with this division of labor, coreceptors IR8a/25a sacrifice ligand-binding competence for the ability to make extensive stabilizing contacts through the beam, reducing the interfaces of and therefore relaxing sequence constraints on tuning receptors. This may enhance mutational robustness of the protein family, supporting the maintenance of large receptor families and permitting more extensive remodeling of tuning-receptor architecture that would otherwise compromise complex stability.

### Structural evolution of subfamily specificity

In coreceptors, we observe ancestral features such as the ligand-binding residues marshalled to structural ends through the gain of novel roles or partners rather than being eliminated: a cautionary tale that sequence conservation does not necessarily imply functional equivalence.

Analogously, among co-receptor paralogs IR8a and IR25a, we observe features conserved in type and position, such as the LBD ion-coordinating site or beam glycosylation, yet engage in variable interactions depending on the tuning receptor. This structural plasticity is precisely the strength of co-receptors, which flexibly accommodate varied interaction modes from diversified tuning receptors.

However, the same structural plasticity likely provided the substrate for the deep evolutionary divergence of IR subfamilies. The ancestral IR25a, which gave rise to IR8a through gene duplication in the common ancestor of arthropods (Croset 2010, Eyun 2017), presumably accommodated multiple modes of tuning-receptor assembly. Following duplication, IR8a co-evolved with a specific monophyletic lineage of tuning receptors, accumulating mutations entrenching this interface and precluding assembly with other tuning lineages. IR25a, in contrast, retains ancestral flexibility, remaining associated with a massive radiation of tuning receptors and secondary co-receptors to sense highly distinct environmental cues.

### Ancestral receptor architecture shapes ligand specificity

The evolutionary repurposing of the iGluR scaffold likely constrains the chemical specificity of IRs. While ORs bind hydrophobic odorants within buried transmembrane pockets, IRs retain the soluble, extracellular LBDs of their iGluR ancestors, suited to detect polar or charged soluble compounds (e.g., acids, amines) via electrostatic complementarity. Yet, receptor architecture does not absolutely dictate ligand chemistry: IRs are also known to detect hydrophobic compounds like cholesterol (Pradhan 2025), and may support sophisticated aerial olfaction in OR-lacking arthropods such as ticks, mites, and coconut crabs (Stensmyr 2005, Josek 2018, Hoy 2016). How they may do so remains an outstanding question.

### Coupled gating mechanisms

While IR tuning subunits diversified, primary co-receptors retained highly conserved M3 hinge regions, suggesting they might actively participate in channel gating. Our structures show novel interactions between gating regions of tuning and co-receptor subunits that could enable a coupled gating mechanism. In that context, the asymmetric conservation of the SYTANLAAF motif could be interpreted considering the differential role of distal and proximal subunits in channel gating. In iGluR tetramers, subunits occupying distal positions feature horizontally splayed M3-D2 linkers, granting them a mechanical advantage during gating compared to the vertical linkers of proximal subunits (Kazi 2014, Amin 2021, Yelshanskaya 2022, Chou 2024, Gangwar 2025). In the IR8a/IR75b structure, the tuning subunits occupy these mechanically advantageous distal positions (Figure 4b). Because of this horizontal geometry, their degenerate M3 helices merely need to be flexible enough to yield to tension when the LBD closes upon ligand binding. Conversely, the co-receptors occupy the proximal positions, where the vertical transmission of tension is mechanically less efficient. Consequently, the co-receptors might rely on the conserved, specialized SYTANLAAF hinge to successfully execute pore opening when acted upon by the tuning subunits.

## Limitations of this study

Unlike IR25a-dependent subunits, IR8a-dependent tuning receptors are relatively recently derived and form a monophyletic clade. Therefore, it is difficult to determine whether differences we point out to define subfamily specificity are really structural requirements for assembly or merely phylogenetically contingent features of this clade. Further assembly and functional assays coupled to mutations or swaps of proposed features will reveal ultimate determinants of subfamily specificity.

We note that although previous studies using heterologous expression in Xenopus oocytes have reported ligand-evoked currents from IR complexes, we have not been able to consistently reproduce these responses. As with many iGluRs that are notoriously reliant on auxiliary components for function, IR function may require additional auxiliary subunits or cofactors. Understanding how gating occurs in the various IR assemblies and how they are remodeled to sense noncanonical hydrophobic ligands or transduce temperature and humidity will require the development of robust functional assays.

## MATERIALS AND METHODS

### Construct design

IR8a and IR75b genes were obtained from a cDNA library generated from antennae of male *Rhodnius prolixus*. Antennae were ground, total RNA was extracted with TRIzol (Invitrogen), and cDNA was synthesized using SuperScript III reverse transcriptase (Invitrogen).

IR8a(Δ809) was generated by deletion of the last 49 amino acids and subsequently fused to mCherry, hemagglutinin (HA), and protein C tags. Full-length IR75b was fused to sfGFP, FLAG, and Strep-tag II. All tags were linked to the respective IR subunit via an HRV 3C protease cleavage site.

Inserts were cloned into a modified pVL1393 transfer vector (pVL1393-WPRE-SV40), in which WPRE and an SV40 polyadenylation signal were inserted downstream of the polyhedrin promoter expression cassette. Coding sequences of *Anopheles gambia*e IR25a (#), IR76b (#) and IR41a (#), *Drosophila melanogaster* IR93a (Genbank #) and *Vanessa cardui* IR40a (#) were codon optimized and synthesized (Twist Bioscience).

For agamIR25a/76b/41a, the genes were cloned into VLBM7 with mCherry, hemagglutinin (HA), and protein C tags for IR25a and IR76b, and sfGFP, FLAG, and Strep-tag II tags for IR41a.

For IR25a/93a/40a, sequences excluding native N-terminal signal peptides were cloned into custom tetracycline -inducible vectors with the influenza hemagglutinin signal peptide followed by an epitope tag (Protein C for IRcos IR25a and IR93a, FLAG for IR40a), and C-terminal mScarlet tags for IRcos and sfGFP for IR40a.

### Expression and purification of IR8a/IR75b, IR25a/IR76b/41a, and IR25a/IR93a/IR40a complexes

#### IR8a/IR75b expression

ExpiSf9 cells (Thermo Fisher) were used both to generate baculoviruses and for protein expression. Cells were grown at 30 °C in ExpiSf™ CD Medium (Thermo Fisher) to a density of 5 × 10^6 cells/mL and transduced with 10% (v/v) baculovirus for each construct. Twenty-four hours post-transduction, ExpiSf enhancer was added according to the manufacturer’s instructions. Cells were harvested by centrifugation 72 hours post-transduction.

#### IR25a/IR76b/IR41a expression

Sf9 insect cells (Expression Systems) were used to generate baculoviruses encoding each construct. HEK293S GnTI− cells (ATCC CRL-3022) were grown at 37 °C with 8% CO₂ in Freestyle 293 medium (Gibco) supplemented with 2% (v/v) fetal bovine serum (Gibco). Cells were grown to a density of 1 × 10^6 cells/mL and transduced with 10% (v/v) baculovirus. After 24 hours, 10 mM sodium butyrate (Sigma-Aldrich) was added and the temperature was shifted from 37 °C to 30 °C for the remainder of the incubation. Cells were harvested by centrifugation 48 hours post-transduction.

#### IR25a/IR93a/IR40a expression

Expi293™ Tetracycline-Inducible cells (Expression Systems) were grown at 37 °C with 8% CO₂ in Freestyle 293 Expi293 (Gibco) medium. Cells were grown to a density of 2.5 × 10^6 cells/mL and transiently transfected at a ratio of 500 ng of plasmid DNA for each subunit mixed with 100 µl of Optimem (Gibco) and 1.2 µl of FectoPRO transfection reagent (Genesee) per mL of cells. After 48 hours, expression was induced with the addition of 4 µl/mL doxycycline, enhanced with 100mM sodium butryate and 0.35% glucose, and moved to 30 °C for an additional 24 hours. Cells were then pelleted by centrifugation, flash frozen, and stored at −80 °C until the day of purification.

#### Purification

Approximately 20 g of frozen cell pellet were thawed and resuspended in 200 mL solubilization buffer containing 50 mM HEPES/NaOH (pH 7.5), 150 mM NaCl, 0.5% (w/v) lauryl maltose neopentyl glycol (LMNG; Anatrace), 0.1% (w/v) cholesterol hemisuccinate (CHS; Sigma-Aldrich), 2.5 µg/mL trypsin inhibitor, 2.5 µg/mL leupeptin, 2.5 µg/mL pepstatin, 5 mM benzamidine, and 1 mM phenylmethylsulfonyl fluoride (PMSF). Cell suspensions were homogenized with a Dounce homogenizer on ice and then stirred with a magnetic stir bar at 4 °C for 1 hour. The lysate was clarified by centrifugation at 175,000 × g for 40 minutes, and the supernatant was incubated with 3 mL anti-GFP nanobody–coupled Sepharose resin.

The resin was washed with buffer containing 50 mM HEPES (pH 7.5), 150 mM NaCl, 0.05% LMNG, and 0.001% CHS, followed by a second wash containing 0.01% LMNG and 0.002% CHS. Bound complexes were eluted by incubating the resin with 25 µg HRV 3C protease (Sigma-Aldrich) per 1 mL resin with rotation at 4 °C for 2 hours.

For IR25a/IR93a/IR40a complexes, 2mM CaCl₂ was added to the clarified supernatant and the mixture incubated with 1 mL M1 Anti-dykddddk (FLAG)-coupled Sepharose resin with rotation at 4 °C for 1 hour. The resin was washed with 20 column volumes of buffer containing 2mM CaCl₂, 50 mM HEPES (pH 7.5), 150 mM NaCl, 0.01% LMNG, and 0.001% CHS, and the complexes were eluted with ∼8 mLs of buffer containing 2 mM EDTA, 100 µg/mL FLAG peptide (Sigma), 50 mM HEPES (pH 7.5), 150 mM NaCl, 0.05% LMNG, 0.001% CHS, and concentrated.

The elution was loaded onto a Superose 6 Increase column (Cytiva) pre-equilibrated with 20 mM HEPES (pH 7.5), 150 mM NaCl, 0.002% LMNG, and 0.0004% CHS. Peak fractions containing IR/IR-co-receptor complexes were pooled and concentrated to 2 µg/µL. Proteins were flash-frozen in liquid nitrogen and stored at −80 °C until use.

#### Cryo-EM sample preparation and data collection

Cryo-EM grids were prepared using a Vitrobot Mark IV (FEI). Three microliters of concentrated protein sample were applied to glow-discharged UltraAuFoil R1.2/1.3 300-mesh gold grids, blotted for 5 s at 100% relative humidity and 4 °C, and plunge-frozen in liquid ethane cooled by liquid nitrogen.

Cryo-EM data were collected on a Titan Krios transmission electron microscope (FEI) operated at 300 kV and equipped with a Gatan BioQuantum GIF and K3 direct electron detector. Automated data collection was performed using SerialEM. Movies were recorded as 50 frames with a total electron dose of 50 e⁻/Å² and a defocus range of −1.5 to −2.0 µm. A portion of the dataset was collected with a 30° stage tilt.

#### Image processing

All cryo-EM data were processed using cryoSPARC v4.7.1. Movie frames were motion-corrected and contrast transfer function (CTF) parameters were estimated in cryoSPARC Live. Particles were initially picked using the blob picker with a particle diameter range of 100–300 Å, and extracted using a box size of 440 pixels. Multiple rounds of 2D classification were performed to enrich tetrameric particle projections. Selected 2D class averages were used as templates for template-based particle picking, followed by an additional round of 2D classification. Good particles were used to generate initial 3D maps using ab initio reconstruction. One of the ab initio volumes served as a reference for subsequent heterogeneous refinement to remove junk particles. After 2D classification and multiple rounds of heterogeneous refinement, a “seed” strategy was employed to enrich good particles from the full particle stack. Heterogeneous refinement was carried out, good classes were combined, and duplicate particles were removed. Several iterations of seed-facilitated heterogeneous refinement and non-uniform refinement were performed. Good particles were then re-extracted and subjected to 3D classification with a mask focusing on the transmembrane domain (TMD). Final processing included non-uniform refinement, reference-based motion correction, and local refinement, resulting in high-quality density maps. The IR8a/IR75b dataset was used to train a Topaz model that was subsequently applied during particle picking for the IR25a/IR76b/IR41a dataset.

#### IR8a/IR75b

A total of 11,077 movies were collected. 3D classification yielded two major classes: one containing the full-length IR8a/IR75b complex with the IR75b ligand-binding domain (LBD) in an open conformation, and another containing a complex in which the IR75b LBD was closed and the TMD was not resolved. After further processing, the full-length complex yielded a high-quality map with an overall resolution of 2.9 Å, while the second class yielded a map at 3.0 Å overall resolution, as estimated in cryoSPARC using the Fourier shell correlation (FSC) 0.143 criterion.

#### IR25a/IR76b/IR41a

A total of 10,444 movies were collected. After processing, only the IR25a/IR76b complex could be reliably identified. The full-length complex yielded a high-quality map with an overall resolution of 2.6 Å, and a second class produced a 3.0 Å map, as estimated in cryoSPARC using the FSC 0.143 criterion.

#### IR25a/IR93a/IR40a

A total of 5,787 movies were collected. 3D classification with a mask on the IR40a LBD yielded multiple classes with the D2 lobe in various positions. The map used in this paper, with the D2 lobe in the most open position, has an overall resolution around 3.0 Å using the FSC 0.143 criterion.

#### Model building, refinement, and structural analysis

The AlphaFold 3 model of the IR8a/IR75b complex was used as an initial model for manual model building in Coot. AlphaFold 3 models of IR25a/IR76b and IR25a/IR93a/IR40a were used as a starting models for manual model building in ISOLDE. The resulting cryo-EM density maps were of sufficient quality to build the majority of the protein chains.

Figures of the maps and atomic models were generated using PyMOL and UCSF ChimeraX.

#### Phylogenetic analysis

We collected iGluR sequences from Ramos-Vicente 2018 and Singh 2025, with particularly dense sampling of AKDF family receptors from bilateria, cnidaria, placozoa, and porifera. These were combined with coreceptor sequences from species across protostomes, as well as insect and C. elegans tuning receptors with known coreceptor partners. We aligned sequences in Jalview using MAFFT L-INS-i (Waterhouse *et al*. 2009, Katoh & Standley 2013) and trimmed in ClipKIT (Steenwyk *et al*. 2020) in default smart-gap mode. We constructed maximum likelihood phylogenetic trees with IQ-TREE v3 (Wong *et al*. 2026), using the site-heterogeneous LG+C60+F+G model. We calculated support for clades with UFBoot (1,000 replicates) and SH-aLRT (1,000 replicates). All sequences used and their sources are collected in Supplementary table 2.

#### Tuning receptor specificity analyses

We compiled a list of reported co-receptor functional requirements for various tuning receptors in the literature. These include *in vivo* evidence (knockout or complementation data), reports of successful reconstitution of heterologous function, as well as findings from our own lab. The information is compiled in Supplementary table 3. Features were evaluated through visual examination of AlphaFold models; publicly unavailable models are indicated in table 3 and included in the supplementary files. To analyze sequence conservation, we aligned the sequences in Jalview using MAFFT L-INS-i. As the ATDs are often poorly conserved between tuning receptors on a sequence level, we additionally extracted the predicted or experimentally determined receptor ATDs used Foldseek (Kim *et al*. 2025) to perform structural alignment (Supplementary files). Visualization of sequence conservation was performed using ConSurf (Celniker *et al*. 2013).

## ACKNOWLEDGEMENTS

We thank Leslie Vosshall, Eric Gouaux, Marie Bao, Richard Benton, Edward Twomey, Wendy A. Valencia-Montoya, and members of the del Mármol lab for discussion and advice on the manuscript. We thank Richard Walsh, Megan Mayer, Conny Leistner and the Structural Biology Center at Harvard and Rui Yan and the Janelia CryoEM facility for help with data collection. We also thank the SBGrid Consortium for data processing resources and support. This article is subject to HHMI’s Open Access to Publications policy. HHMI lab heads have previously granted a nonexclusive CC BY 4.0 license to the public and a sublicensable license to HHMI in their research articles. Pursuant to those licenses, the author-accepted manuscript of this article can be made freely available under a CC BY4.0 license immediately upon publication.

## AUTHOR CONTRIBUTIONS

J.dM. G.D.P and A.Q.C conceptualized and supervised the project. G.D.P, A.Q.C and M.S. performed protein expression, purification and specimen preparation for cryo-EM. G.D.P, A.Q.C, J.Z. collected the cryo-EM data. G.D.P and J.Z. built the molecular models. G.D.P, A.Q.C. J.Z. J.R. analyzed the molecular models. A.Q.C performed alignments and phylogenetic analyses. J.M.L.E and M.L. provided R. prolixus antennae samples, processing methodology and contributed to data interpretation. J.I.dM. G.D.P A.Q.C wrote the manuscript, which was then edited by all authors.

## SUPPLEMENTARY FIGURES

**Supp figure 1.**
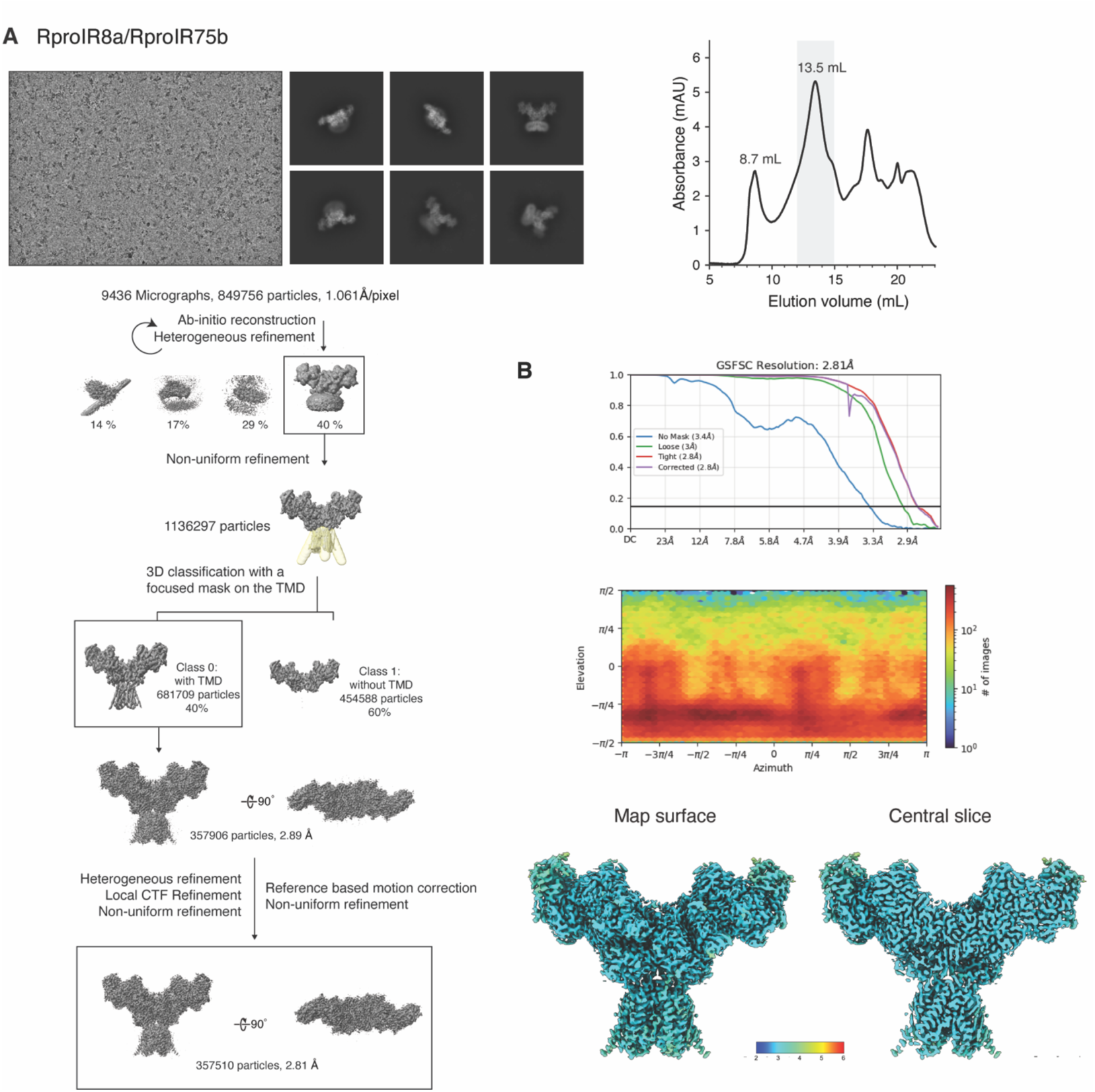
Protein purification and cryo-EM data analysis of the RproIR8a/IR75b dataset. (A) Right: Size-exclusion chromatography profile for purification of IR8a/75b, with protein A280 absorbance plotted against elution volume from column. Fractions indicated in grey were pooled and concentrated for Cryo-EM. Left top: representative micrographs for IR8a/75b. Below: Cryo-EM processing workflow for IR8a/75b structure. (B) GFSC local resolution and particle orientations for IR8a/75b.

**Supp figure 2.**
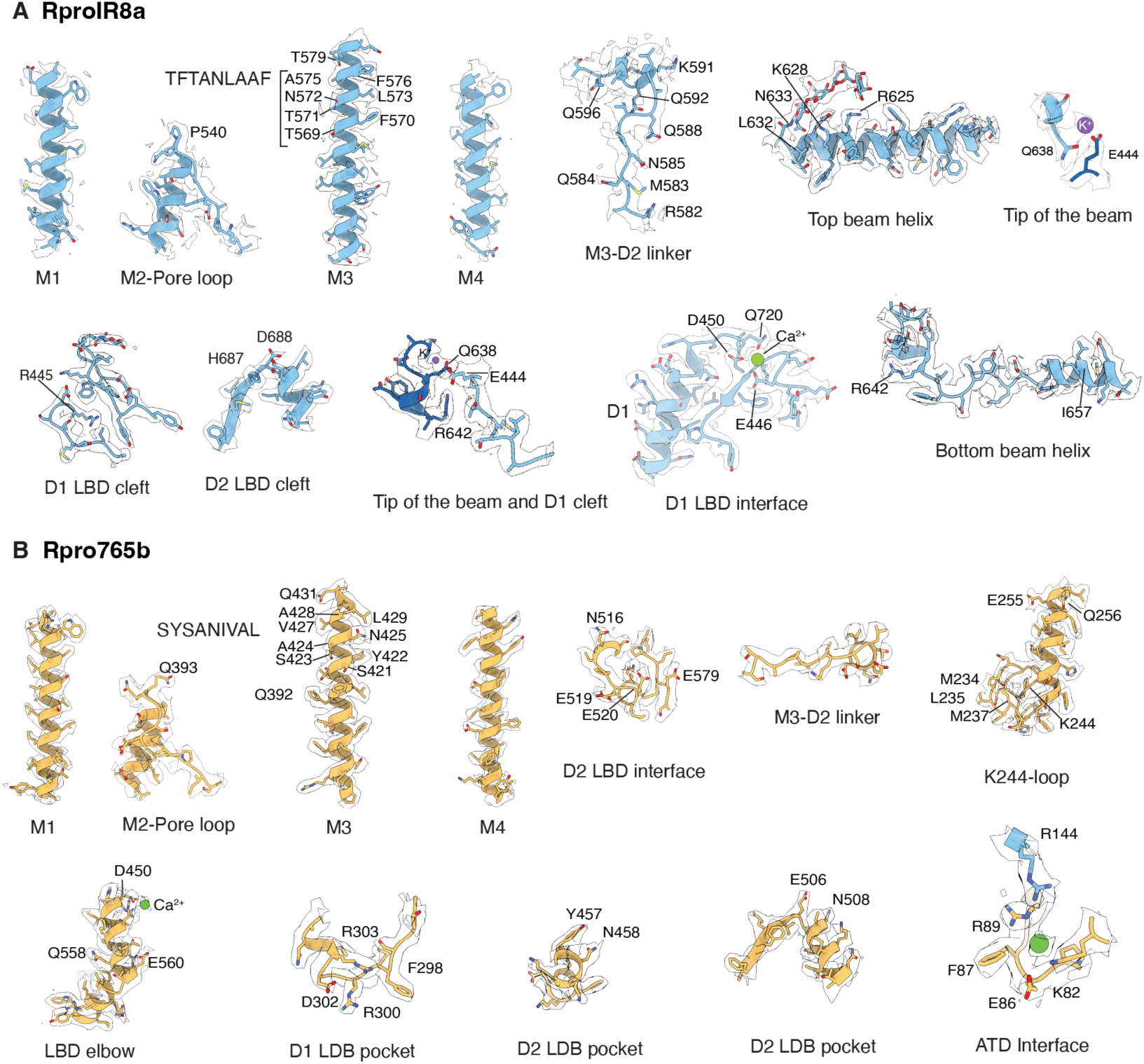
Details of the cryo-EM map of RproIR8a/RproIR75b. (A) RproIR8a density and model for the M1-M4 transmembrane helices, the M3-D2 linker, the top and bottom beam helices, the beam tip, and the D1 and D2 LBD clefts and interfaces. (B) RproIR75b density and model for the M1-M4 transmembrane helices, the M3-D2 linker, the LBD elbow, the K244 loop, the D1 and D2 LBD clefts, and the ATD interface.

**Supp figure 3.**
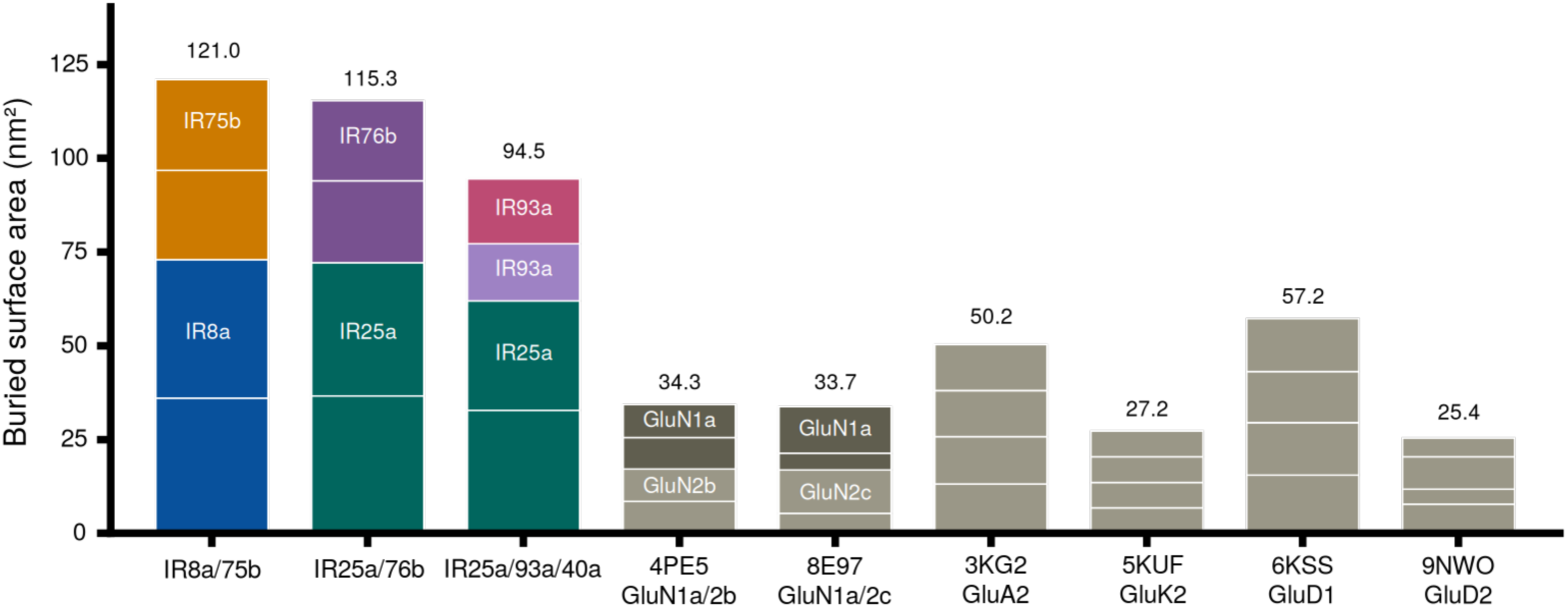
Comparison of LBD buried surface areas (BSA) across different IR/iGluR assemblies. In each column, boxes represent the relative contribution of each subunit to the total BSA. PDBs are indicated for published iGluR structures.

**Supp figure 4.**
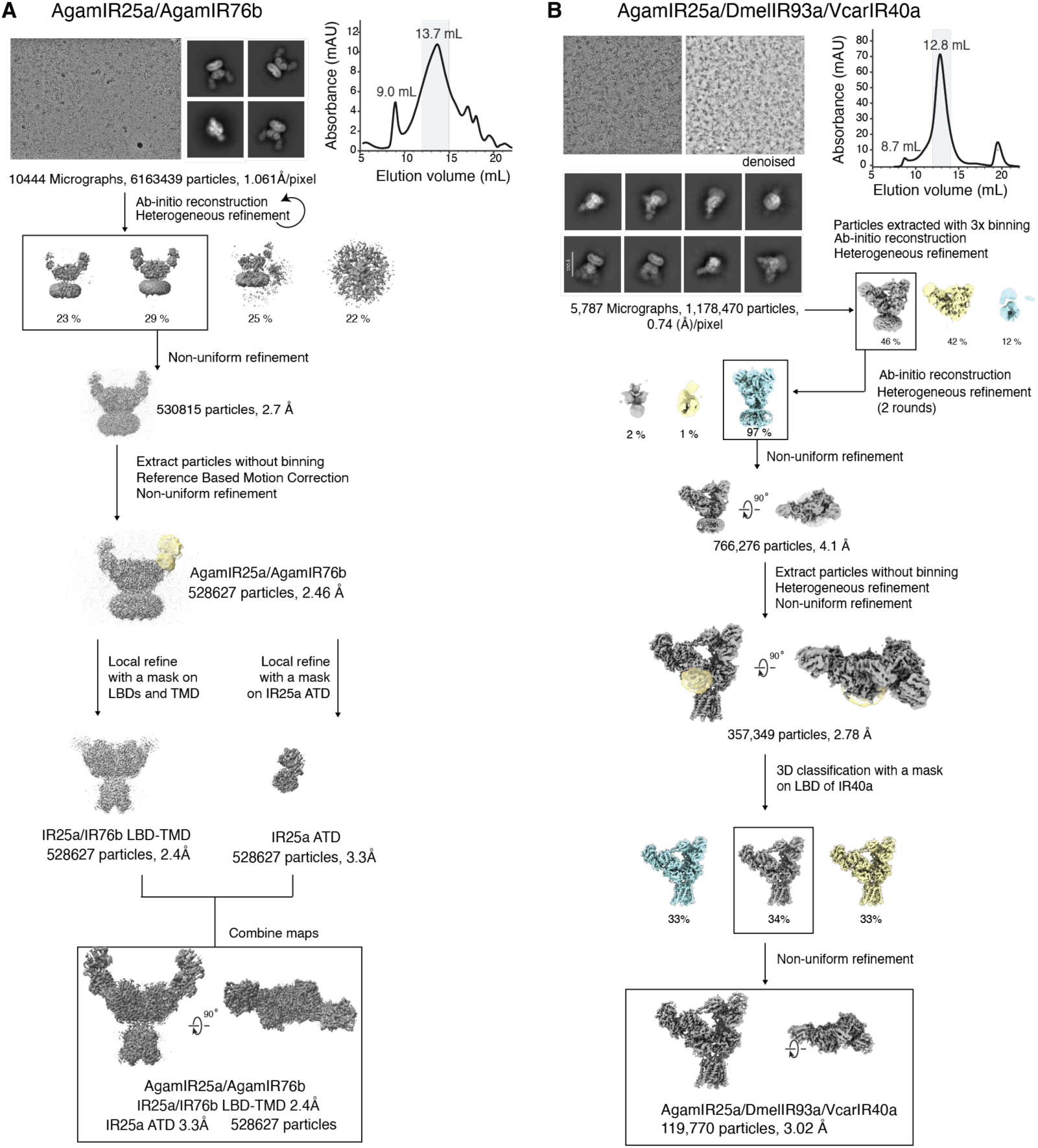
Protein purification and cryo-EM data analysis of the AgamIR25a/AgamIR76b (A) and AgamIR25a/DmelIR93a/VcarIR40a datasets (B). For each dataset: representative micrograph and 2D class averages (left), size-exclusion chromatography profile with pooled fractions indicated in grey (right), and the cryo-EM processing workflow from ab-initio reconstruction and heterogeneous refinement through focused 3D classification and non-uniform refinement to the final reconstructions, with particle numbers and resolutions indicated at each step.

**Supp figure 5.**
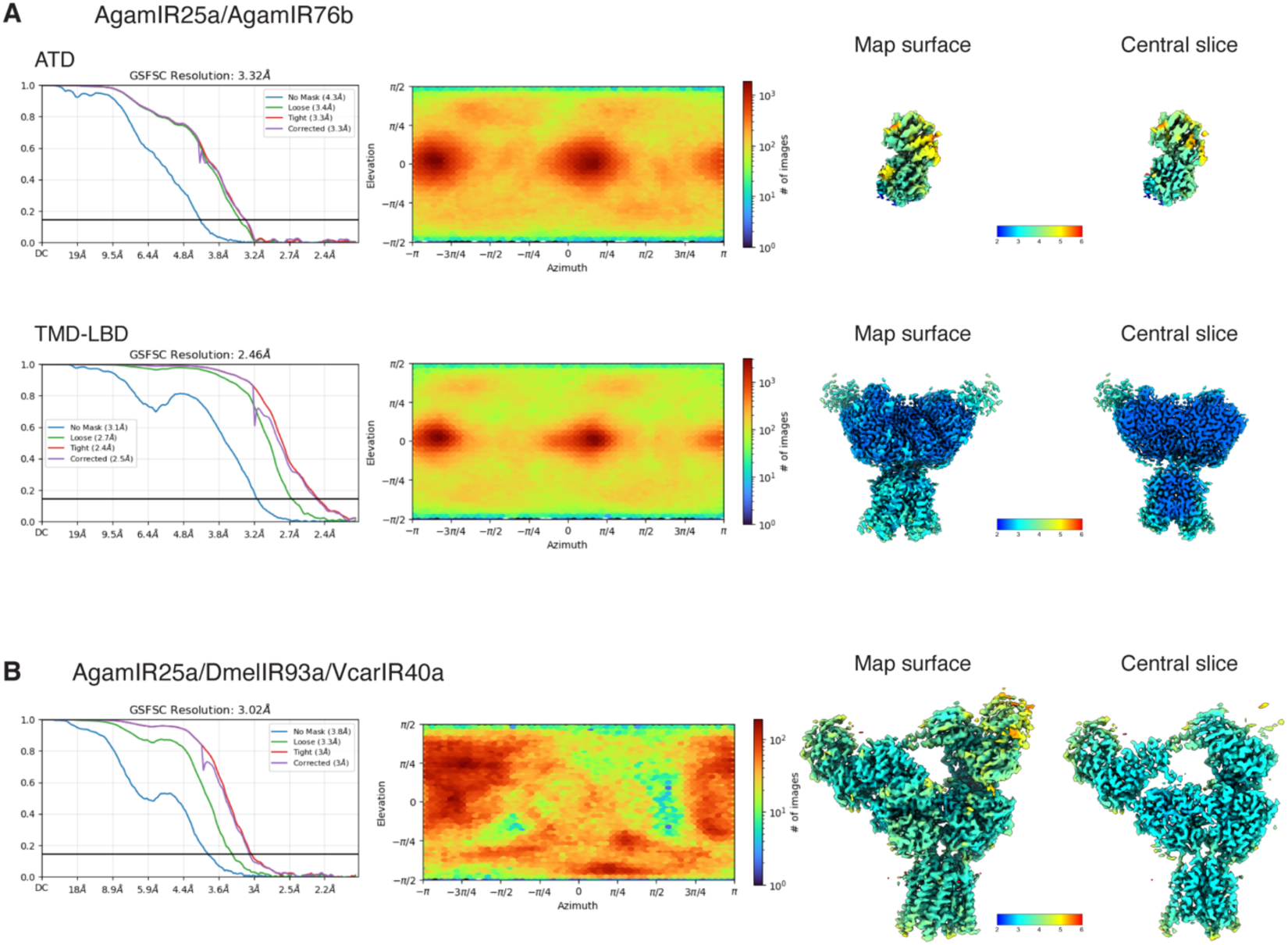
GSFSC local resolution and particle orientations for AgamIR25a/AgamIR76b (A) and AgamIR25a/DmelIR93a/VcarIR40a (B) datasets. For each map (ATD and TMD-LBD reconstructions of AgamIR25a/AgamIR76b in A; consensus reconstruction of AgamIR25a/DmelIR93a/VcarIR40a in B): gold-standard Fourier shell correlation (GSFSC) curves with the indicated resolutions, particle orientation distribution, and the final map colored by local resolution, shown as a surface and as a central slice.

**Supp figure 6.**
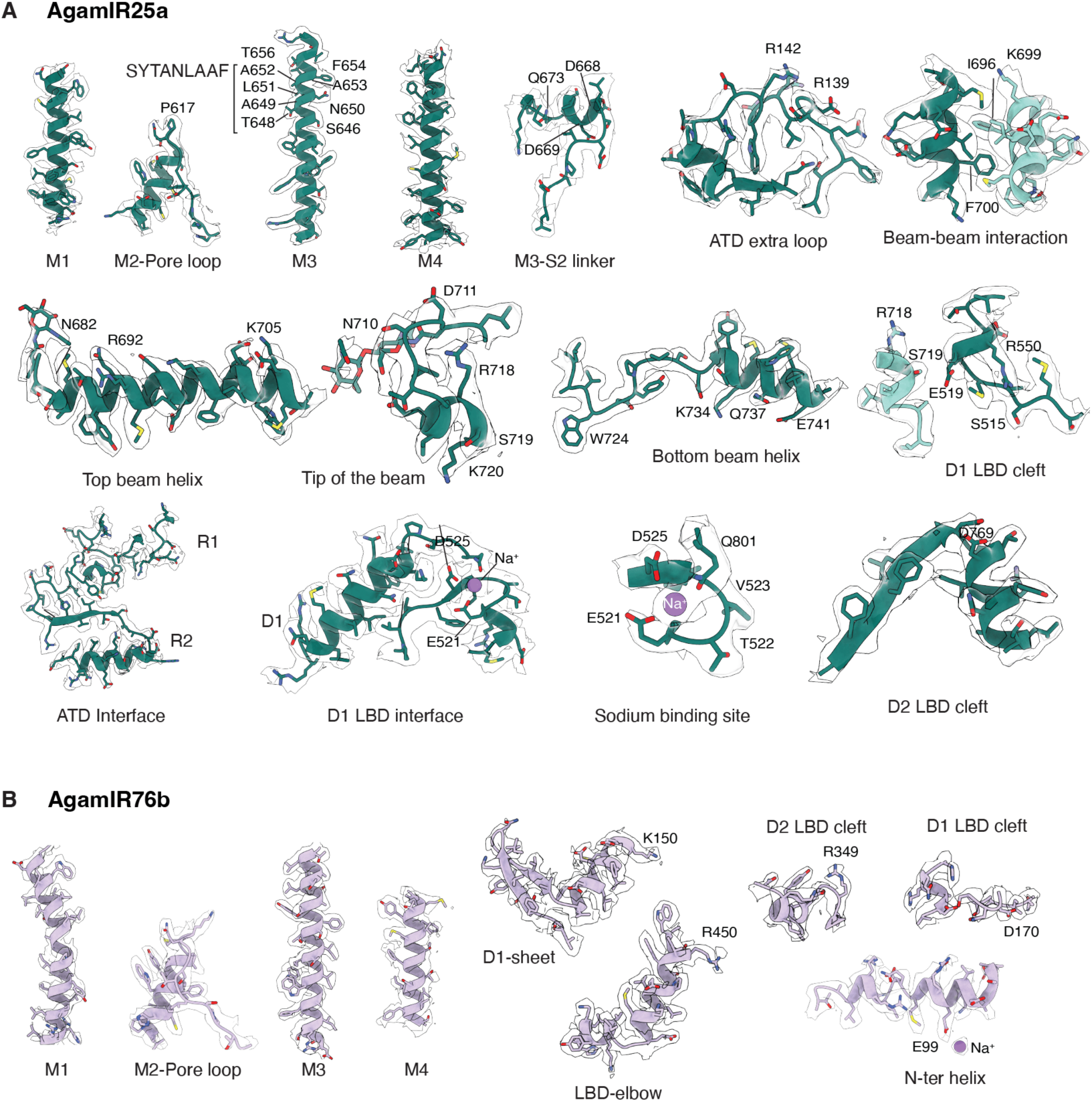
Details of the cryo-EM map of AgamIR25a/AgamIR76b. (A) AgamIR25a density and model for the M1-M4 transmembrane helices, the M3-D2 linker, the ATD extra loop, the beam-beam interaction, the top and bottom beam helices, the ATD and D1 LBD interfaces, the sodium binding site, and the D1 and D2 LBD clefts. (B) AgamIR76b density and model for the M1-M4 transmembrane helices, the D1 beta-sheet, the LBD elbow, the N-terminal helix, and the D1 and D2 LBD clefts.

**Supp figure 7.**
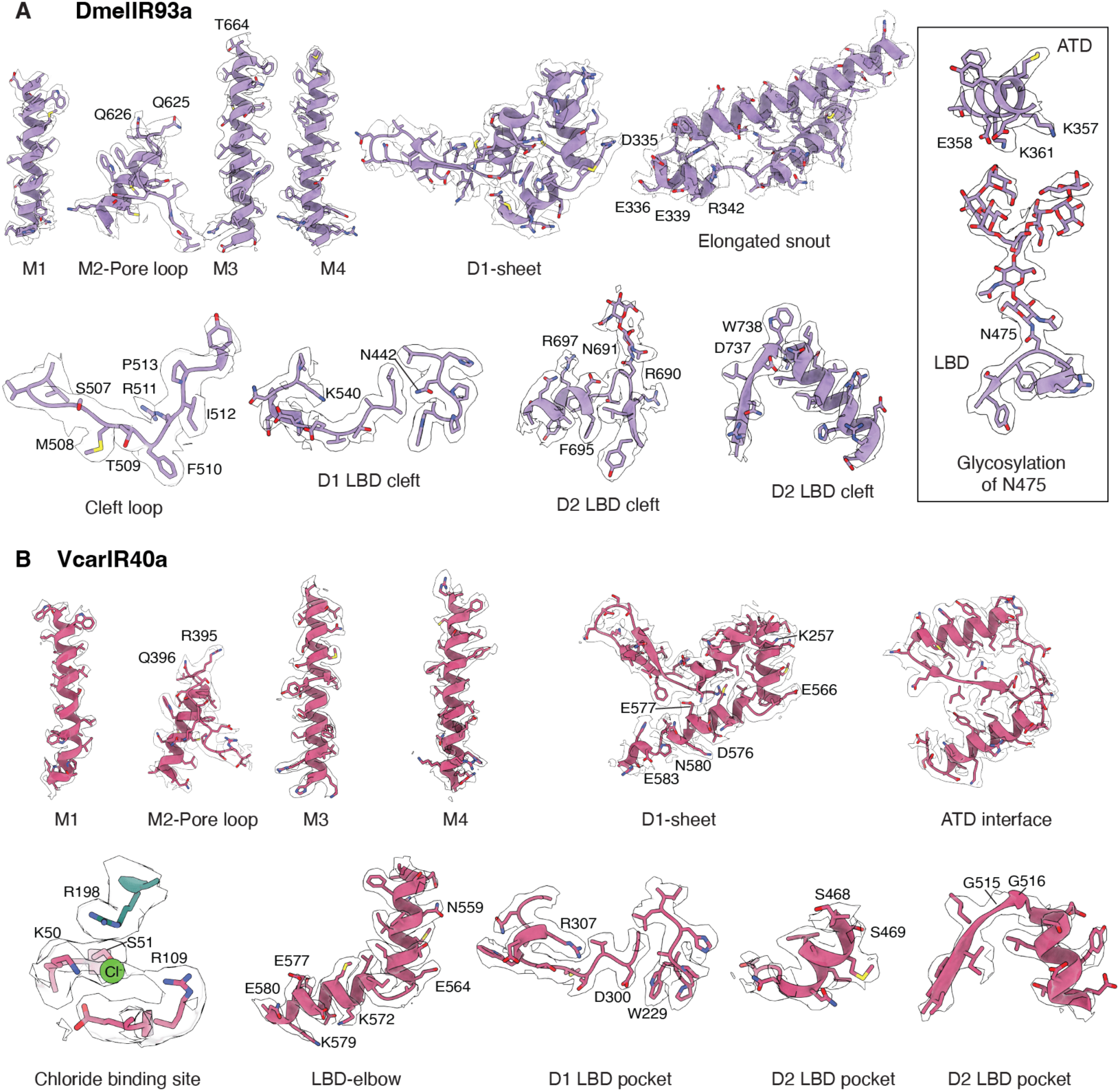
Details of the cryo-EM map of the DmelIR93a and VcarIR40a subunits. (A) DmelIR93a density and model for the M1-M4 transmembrane helices, the D1 beta-sheet, the elongated ATD snout, the cleft loop, the D1 and D2 LBD clefts, and the N475 glycosylation site. (B) VcarIR40a density and model for the M1-M4 transmembrane helices, the D1 beta-sheet, the ATD interface and chloride binding site, the LBD elbow, and the D1 and D2 LBD pockets.

**Supp figure 8.**
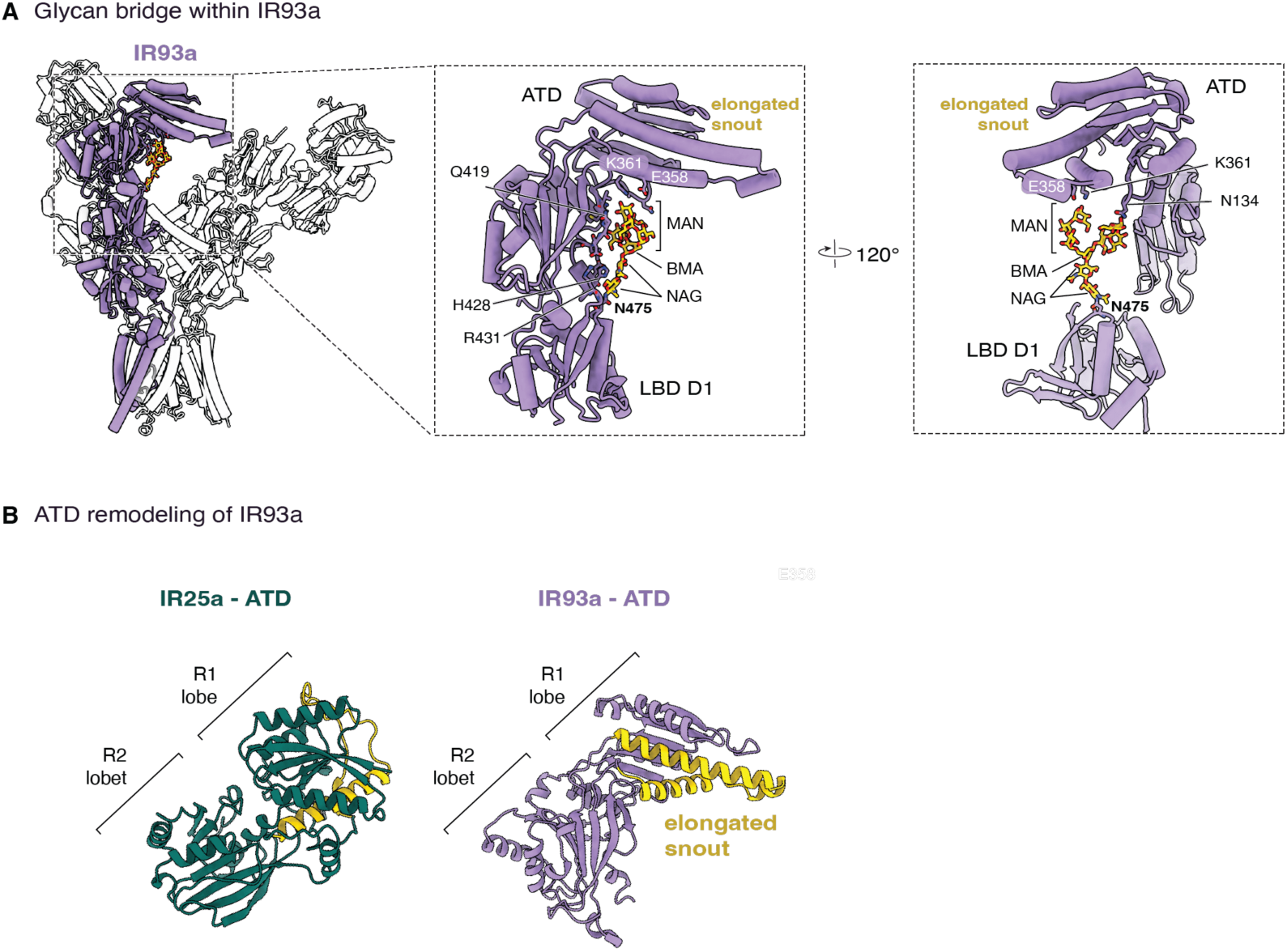
ATD remodeling of IR93a. (A) An elaborate glycan extends from the D1 lobe of the IR93a LBD (at N475) and connects to its own ATD. Cryo-EM density for this post-translational modification is well resolved and allowed building of the entire glycan chain. (B) Comparison of the IR25a ATD (left) and the IR93a ATD (right), highlighting the elongated helical ‘snout’ (yellow) that is unique to IR93a and mediates its bridging interaction with the IR40a ATD.

**Supp figure 9.**
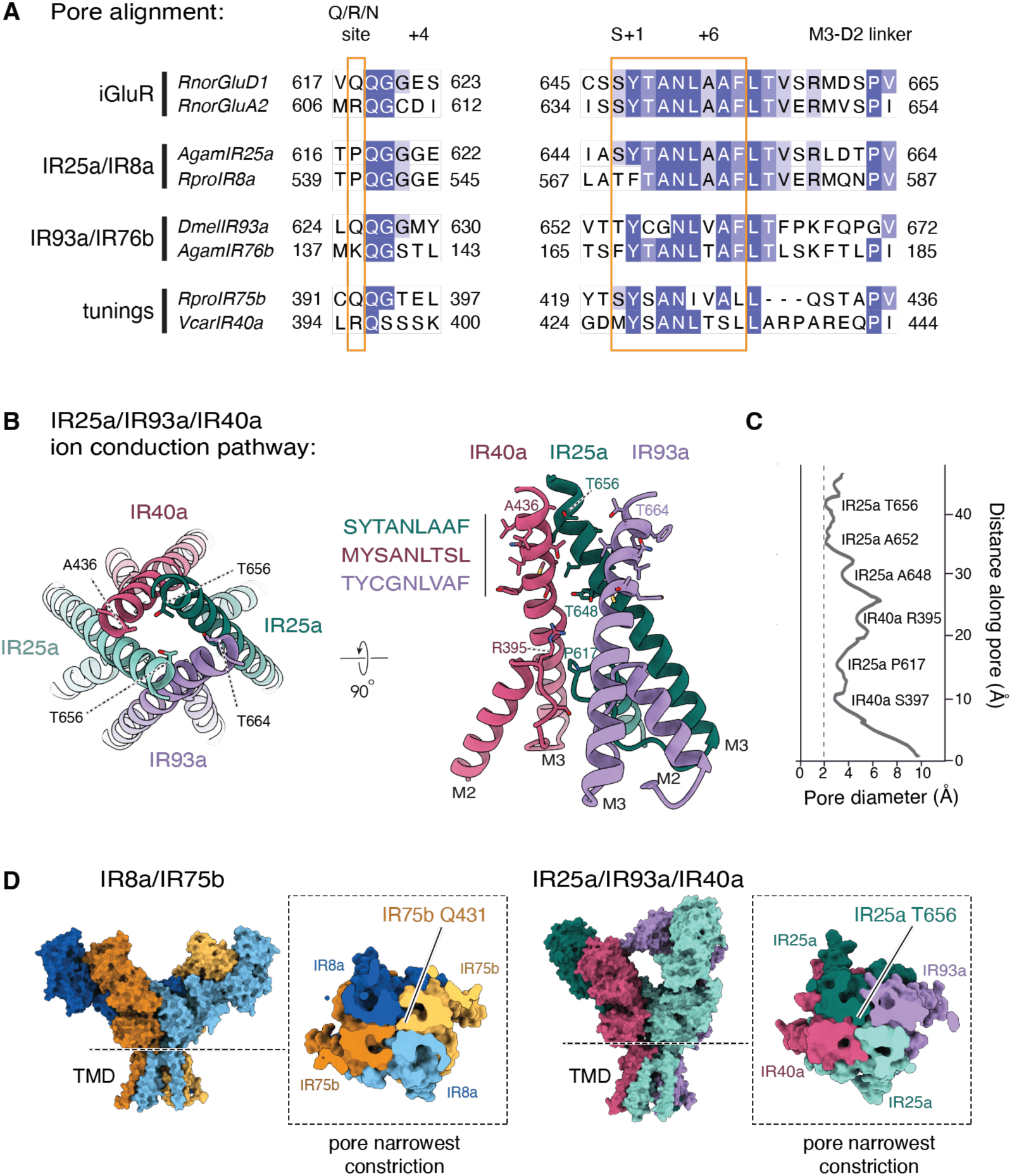
Details of the ion conduction pathway of AgamIR25a/Dmel93a/Vcar40a. (A) Left, alignment of pore loop selectivity filter sequence in representative IRs and iGluRs. The Q/R/N site decisive for calcium permeability in canonical iGluRs is highlighted. Right, alignment of the SYTANLAAF “hinge” area (highlighted) at the end of M3 and the subsequent M3-S2 linker. (B) Left, top view of the pore of IR25a/93a/40a, with M2 and M3 helices shown. M3 residues forming the greatest constriction in each subunit are shown. Right, side view of M2-M3 helices in IR25a/93a/40a, showing constricting residues at the M3 gate and pore loop Q/R/N selectivity filter equivalent residues. (C) Radius (Å) of the AgamIR25a/DmelIR93a/VcarIR40a ion permeation pathway plotted as a function of distance from the extracellular membrane boundary toward the intracellular space. (D) Comparison of the narrowest point along the pore of the AgamIR8a/AgamIR75b and the AgamIR25a/DmelIR93a/VcarIR40a. Surface representation of both complexes in a side view and a slice, showing that both complexes constrict in a pinched configuration, with either the tuning subunits (IR8a complex) or the co-receptor subunits (IR25a complex) closing the pore.

**Supp figure 10.**
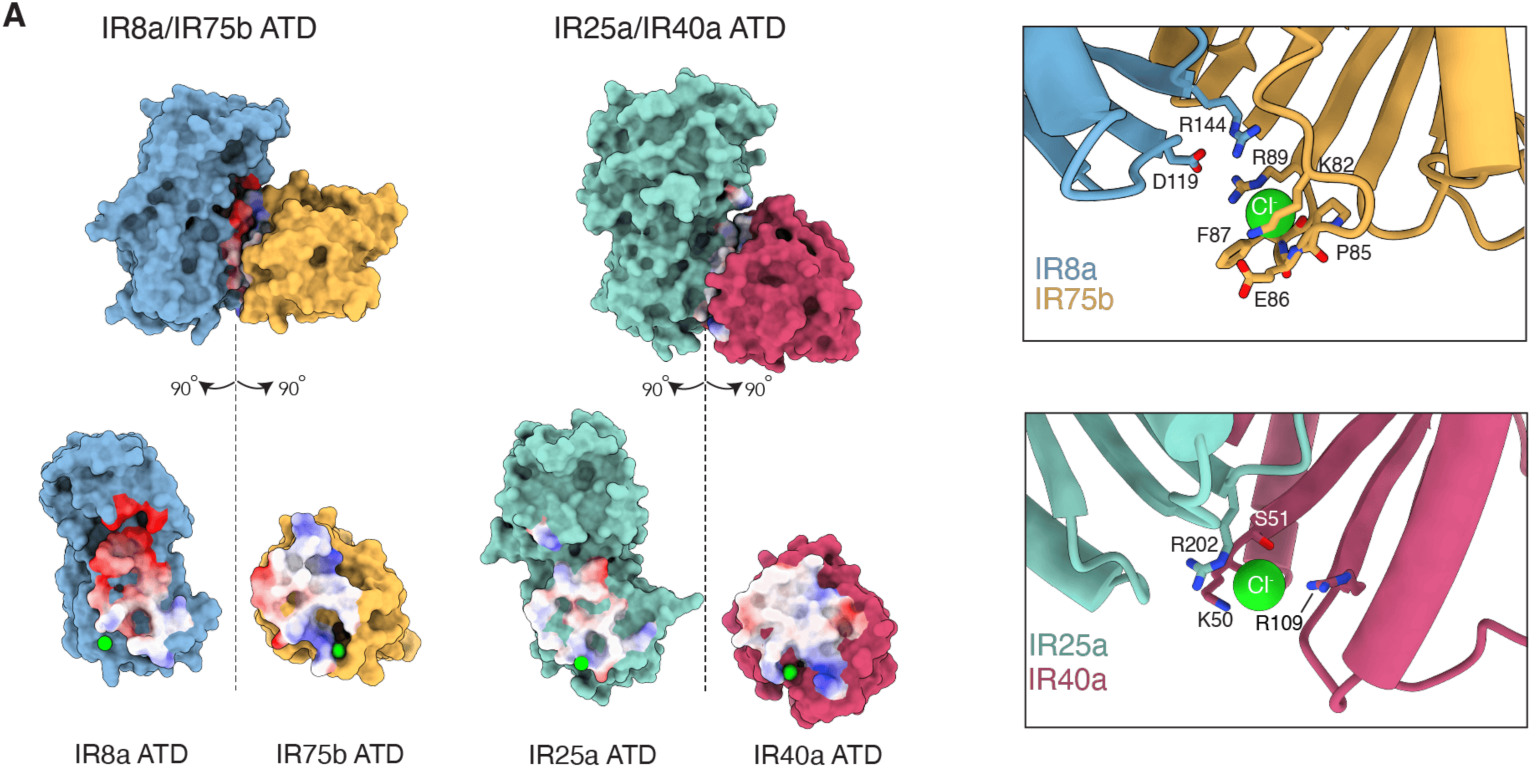
Electrostatic inversion at the ATD level. Top, surface representation of the IR8a/IR75b (left) and IR25a/IR40a (right) ATD pairs. Bottom, electrostatic surface potential of the same interfaces, showing that IR8a presents an electronegative surface that recruits positive charges from the tuning subunit (IR75b), a charge bias not observed at the analogous IR25a-IR40a interface, where the extra loop occludes the cleft and complements its own charged residues. Right, close-up views of the chloride binding site (green) at each ATD-ATD interface, with coordinating residues indicated.

## Supplementary Tables

Supplementary Table 1. Cryo-EM data collection parameters.

Supplementary Table 2. Sequences included in alignment used to construct phylogeny in Figure 2a.

Supplementary Table 3. Curated sequences included in Figure 7 used to compare IR25a- and IR8a-dependent tuning receptors.

